# PTEN regulates starburst amacrine cell dendrite morphology during development

**DOI:** 10.1101/2025.05.08.652956

**Authors:** Teva W. Bracha, Nina Luong, Joseph Leffler, Benjamin Sivyer, Kevin M. Wright

**Affiliations:** Neuroscience Graduate Program, Oregon Health & Science University, Portland, Oregon 97239; Vollum Institute, Oregon Health & Science University, Portland, Oregon 97239; Casey Eye Institute, Oregon Health & Science University, Portland, Oregon 97239

## Abstract

Neurons are subject to extensive developmental regulation to ensure precise subtype-specific morphologies that are intimately tied to their function. Starburst amacrine cells (SACs) in the mammalian retina have a highly stereotyped, radially symmetric dendritic arbor that is essential for their role in direction-selective circuits in the retina. We show that PTEN, the primary negative regulator of the PI3K-AKT-mTOR pathway that is highly implicated in neurodevelopmental disorders, regulates SAC morphology in a cell-autonomous manner.

*Pten*-deficient SACs show a nearly twofold increase in the number of dendritic branches, while other morphological properties remain largely unchanged. These morphological changes arise late in SAC development after dendrite development is largely complete and persist into adulthood. Mechanistically, excessive dendritic branching appears to arise from dysregulated mTOR activity. Despite this dramatic increase in dendritic branches, *Pten*-deficient SACs maintain a normal population number, organization of synaptic outputs, and intact direction-selectivity in the retina. Collectively, these results show that PTEN is essential for the normal development of highly stereotyped neuronal morphology.

## INTRODUCTION

Since the time of Ramón y Cajal, neuroscientists have appreciated the complexity of the nervous system and the vast array of neuronal shapes and sizes. This morphological diversity underlies the computational power of the nervous system, as neurons acquire specific morphologies that are uniquely adapted to support their function [1]. Specific morphological features can be used in conjunction with molecular profiles and functional properties to classify neurons into discrete subtypes [2]. Neuronal morphology is influenced by the interplay between intrinsic factors and extrinsic cues in the extracellular environment. Cell type-specific transcription factors control the expression of effectors that in turn regulate the morphological development of a given neuronal subtype [3]. A neuron’s complement of cell surface receptors allows it to organize its dendritic arbors and identify synaptic partners in response to extrinsic cues. These cell surface receptors converge on intracellular signaling cascades that modulate cytoskeletal dynamics, leading to differences in neurite elongation, branch initiation, and stabilization [4].

Many insights about the development of neuronal morphology come from highly stereotyped neuronal subtypes in a wide range of model organisms. PVD sensory neurons in *C. elegans* have a defined pitchfork-like projection pattern that line the body wall and are fundamental to mechanosensation and proprioception [5]. Forward genetic screens have identified transcription factors, cell surface receptors, and intracellular signaling molecules that are required for the stereotyped PVD neuron projection pattern [6–9]. The dendritic arborization (da) neurons in *Drosophila* larvae can be easily distinguished into four morphologically distinct subtypes based on the degree of dendrite branching [10]. The morphological complexity of the four da neuron subtypes is determined by relative levels of three transcription factors: abrupt, cut, and knot [11–13]. Purkinje neurons in the mammalian cerebellar cortex have large, planar dendritic arbors with extensive branching patterns that maintain a high degree of self-avoidance. Multiple molecular pathways govern the development of these arbors, including repulsive Slit/Robo signaling, protocadherin-mediated self-avoidance, and actin regulators Daam1 and MTSS [14–16].

Starburst amacrine cells (SACs) in the mammalian retina are an excellent model for studying the development of neuronal morphology due to their stereotyped radially symmetric branching pattern, defined circuit function, and the established link between their dendritic form and neuronal function [17, 18]. SAC somas reside in two neuronal layers in the retina, the inner nuclear layer (INL) and ganglion cell layer (GCL), and project their dendrites to the inner plexiform layer (IPL), where they form planar dendritic arbors that stratify in sublamina 2 (S2) and 4 (S4), respectively (Figure 1A). Over the past several years, work from multiple labs has identified cell surface receptors that direct SAC morphology and stratification in the IPL. MEGF10 regulates the mosaic spacing of SACs through mediation of homotypic contacts during development [19–21]. Repulsive signaling mediated by FLRT2/UNC5 regulates SAC dendrite stratification [22]. Bidirectional PlexinA2/Semaphorin6A signaling is critical for SAC radial morphology and dendrite stratification [23, 24]. Other studies have identified molecules that disrupt SAC dendrite morphology without affecting stratification, suggesting that these are separate processes. γ-protocadherins (γ-Pcdhs) undergo extensive alternative splicing to generate hundreds of isoforms, with homophilic matching between isoforms mediating self-recognition in SAC dendrites. Genetic deletion of all *γ-Pcdh* isoforms causes SAC dendrites to fasciculate into bundles, disrupting their radial morphology; expression of a single *γ-Pcdh* isoform in *γ-Pcdh* deficient SACs is sufficient to restore self-recognition and normal radial morphology [15]. Loss of the cell surface protein AMIGO2 results in a 1.5-fold increase in the size of SAC dendritic arbors, but does not affect their stereotyped branching, symmetry, or stratification [25]. While these cell surface proteins are critical for regulating SAC morphology, little is known about the downstream intracellular signaling pathways that govern SAC morphology.

**Figure 1.**
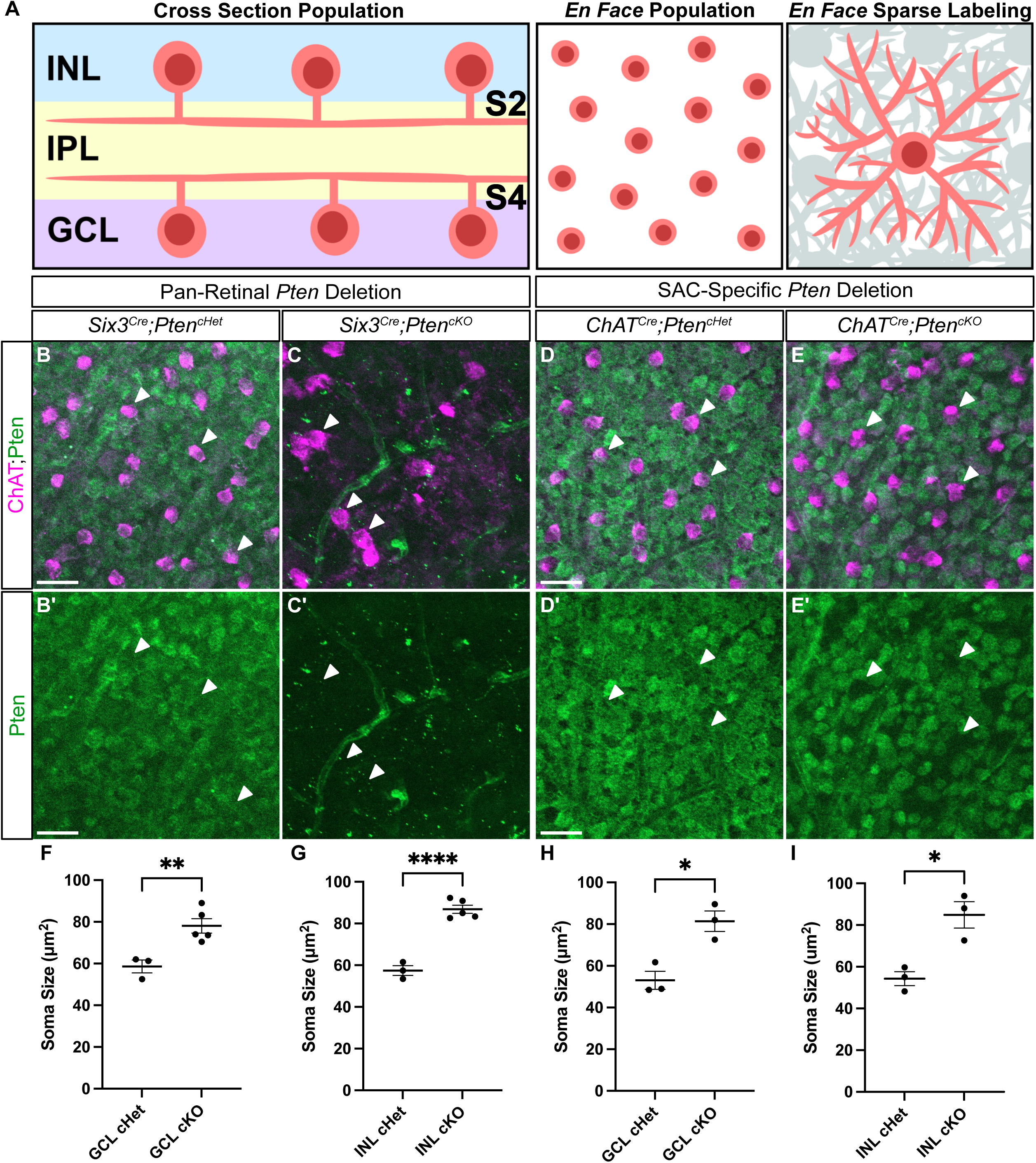
Validation of SAC specific *Pten* deletion. **A**.Schematic showing different retinal preparations for visualizing SACs. Retinal cross-sections (left panel) are used to analyze cellular lamination and dendrite stratification. Retinal flat mounts imaged in an *En Face* preparation are used for population measurements (middle panel) and single cell morphology (right panel). **B-E’.** P28 retinal flat mounts immunostained with ChAT (magenta) to label SAC somas and PTEN (green) shows that PTEN is present in all cells in the GCL. In *Six3^Cre^;Pten^cKO^* retinas, PTEN is eliminated from all GCL cells (C, C’), whereas in *ChAT^Cre^;Pten^cKO^* retinas, PTEN is selectively lost only from SACs (white arrows) (E, E’). **F-I.** Quantification of SAC soma sizes at P28 reveals somal hypertrophy, a common phenotype seen after *Pten* deletion, in both *Six3^Cre^;Pten^cKO^* and *ChAT^Cre^;Pten^cKO^* GCL and INL SACs (p = 0.0433 (F), 0.0017 (G), 0.0124 (H), 0.0307 (I)). Data reported as mean ± SEM. Scale bars = 25 μm.

PTEN (phosphatase and tensin homologue) is a protein and lipid phosphatase that canonically functions as the primary negative regulator of the PI3K-AKT-mTOR pathway [26]. This pathway functions downstream of several cell surface receptors to regulate neuronal differentiation, migration, neurite outgrowth, and survival [27]. PTEN has a well-established role in regulating neurite growth and branching in mammalian neurons *in vivo*. Deletion of *Pten* results in neuronal hypertrophy and increased dendrite branching in cortical pyramidal neurons, hippocampal dentate granule cells, serotonergic raphe neurons, and cerebellar Purkinje neurons [28–31]. This can ultimately lead to altered synaptic connectivity and neuronal hyperexcitability [32, 33].

Whether PTEN plays a role in regulating the highly stereotyped dendritic morphology of SACs remains an open question. Deletion of *Pten* from retinal progenitors results in widespread defects in neuronal differentiation, migration, cellular lamination, mosaic spacing, and dendrite stratification throughout the retina, precluding any analysis of SAC morphology [34–37]. We therefore used a *ChAT^Cre^* line to delete *Pten* specifically from post-migratory SACs (*ChAT^Cre^;Pten^cKO^*) to address its cell-autonomous role in regulating SAC morphology. SACs in *ChAT^Cre^;Pten^cKO^* mice had a >1.5-fold increase in dendritic branching without affecting the overall length or field area of their dendritic arbors. We found that these branching phenotypes arose gradually during the later phase of SAC dendrite development and persist into adulthood. Analysis of signaling pathways downstream of PI3K-AKT suggests that increased dendrite branching is likely due to increased mTOR activity. Finally, we show that loss of *Pten* does not affect the compartmentalization of synaptic outputs in SACs or the function of the direction-selective circuit in the retina.

## RESULTS

### SAC-specific deletion of *Pten* does not affect cell density, somal lamination, mosaic spacing, or dendrite stratification

Pan-retinal deletion of *Pten* from retinal progenitors causes widespread abnormal somal lamination, mosaic spacing, and dendrite stratification in retinal neurons, including SACs [34, 35]. Subsequent work identified a role for PTEN in regulating the vesicular trafficking of cell adhesion molecules that are involved in establishing retinal neuron mosaics and dendrite stratification [37]. However, it is unclear whether these defects reflect a SAC-autonomous effect or are due to the overall disorganization of the retina. To circumvent this confound, we used a *ChAT^Cre^* line to selectively drive recombination in SACs, the only cholinergic neurons in the retina, beginning at postnatal day 1 (P1) [20]. This timing coincides with the end of SAC laminar migration and the initiation of their dendritic stratification in the nascent IPL.

To better understand which aspects of SAC development require cell-autonomous PTEN function, we conducted a side-by-side comparison of pan-retinal (*Six3^Cre^*) and SAC-specific (*ChAT^Cre^*) *Pten* conditional knockouts. Using retinal flat mount preparations from P28 *Six3^Cre^;Pten^cHet^* and *Six3^Cre^;Pten^cKO^* mice, we confirmed that while all cells in the GCL were positive for PTEN in *Six3^Cre^;Pten^cHet^* retinas, staining was completely absent in *Six3^Cre^;Pten^cKO^* retinas (Figure 1B-C’). We next analyzed P28 *ChAT^Cre^;Pten^cHet^* and *ChAT^Cre^;Pten^cKO^* retinas (Figure 1D-E’). In contrast to the complete loss of PTEN staining in pan-retinal mutants, the PTEN staining was selectively lost from ChAT^+^ SAC somas but retained in all other GCL neurons in *ChAT^Cre^;Pten^cKO^* retinas (Figure 1D-E’). As an additional confirmation of functional PTEN loss from SACs, we measured soma size, as neuronal hypertrophy is consistently seen after *Pten* deletion. In both *Six3^Cre^;Pten^cKO^* and *ChAT^Cre^;Pten^cKO^* retinas SAC soma sizes were significantly increased compared to their respective controls (Figure 1F-I).

We next compared the early developmental processes of differentiation, migration, and mosaic spacing in *Six3^Cre^;Pten^cKO^* and *ChAT^Cre^;Pten^cKO^;Ai9* retinas. Since loss of a single *Pten* allele can affect neuronal differentiation in certain contexts, we included both wildtype and heterozygous controls [38–40]. Consistent with previous studies using pan-retinal deletion of *Pten*, we found reduced cellular density and mosaic regularity of SACs in *Six3^Cre^;Pten^cKO^* retinas compared to controls (Figure 2A-E) [34, 35]. In contrast, there was no difference in cellular density or mosaic spacing of SACs in either the GCL or INL following *ChAT^Cre^*-mediated *Pten* deletion (Figure 2F-J). The lack of differentiation or migration phenotypes in *ChAT^Cre^;Pten^cKO^;Ai9* retinas in which SACs are genetically labeled with tdTomato is likely due to *Pten* deletion occurring after these developmental processes are nearly complete and allows us to examine its cell-intrinsic role during dendrite development without these confounds.

**Figure 2.**
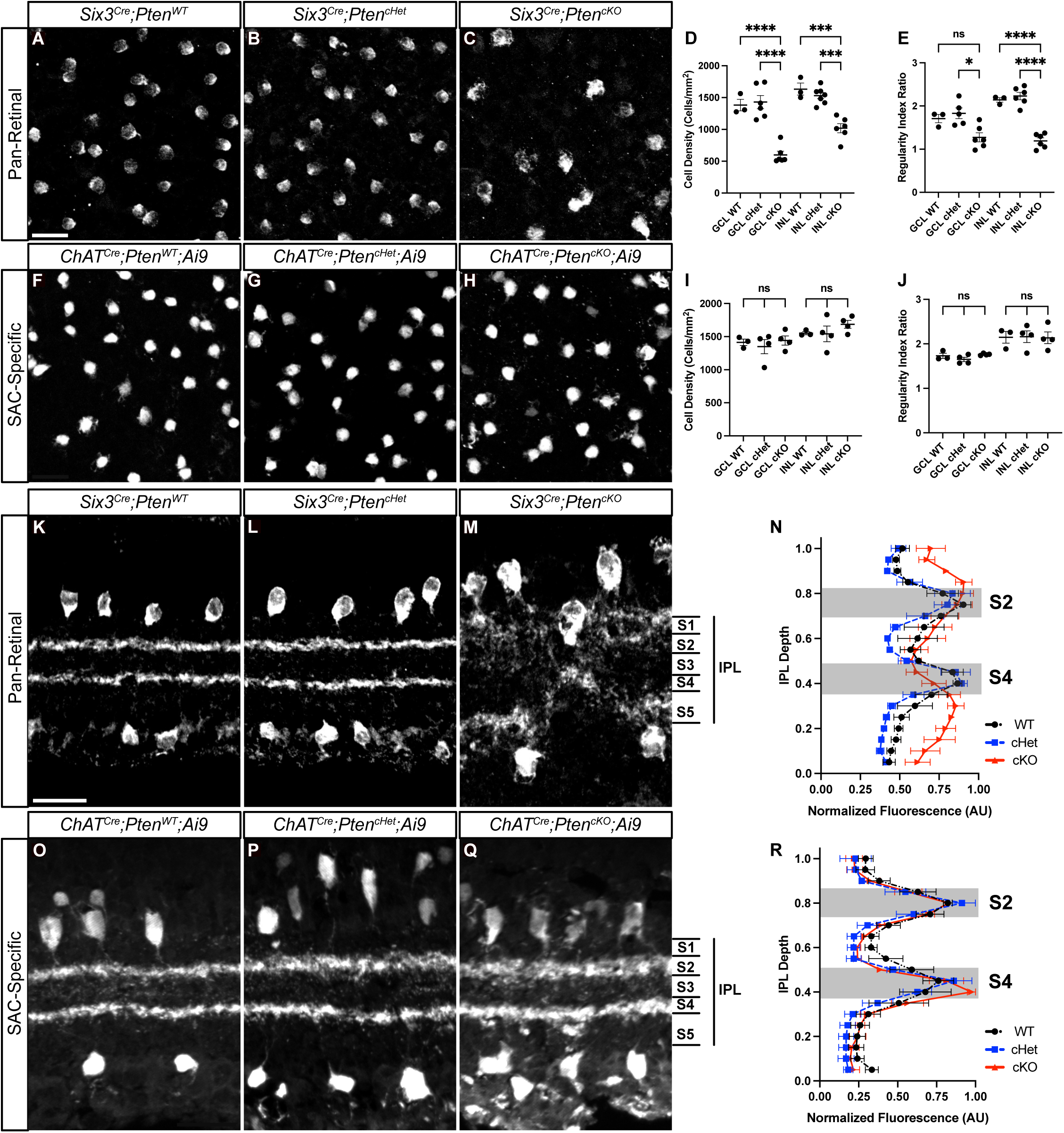
Selective deletion of *Pten* from SACs does not affect their cell density, mosaic spacing, or dendrite lamination. **A-C**. Images of P28 *Six3^Cre^;Pten^WT^*, *Six3^Cre^;Pten^cHet^* and *Six3^Cre^;Pten^cKO^* retina flat mounts with GCL SACs labeled by ChAT immunostaining. **D, E.** Quantification of cell density and mosaic spacing of GCL and INL SACs shows decreased cell density and mosaic regularity in *Six3^Cre^;Pten^cKO^* retinas following pan-retinal deletion of *Pten* (p < 0.0001 for GCL and INL (D), p = 0.01 for GCL and p < 0.0001 for INL (E)). **F-H.** Images of P28 *ChAT^Cre^;Pten^WT^;Ai9*, *ChAT^Cre^;Pten^cHet^;Ai9* and *ChAT^Cre^;Pten^cKO^;Ai9* retina flat mounts with GCL SACs labeled by tdTomato. **I, J.** Quantification shows normal cell density and mosaic spacing of SACs in *ChAT^Cre^;Pten^cKO^;Ai9* retinas following selective deletion of *Pten* from SACs (p = 0.7195 for GCL and 0.4336 INL (I), p = 0.1871 for GCL and 0.9901 for INL (J)). **K-M.** P28 *Six3^Cre^;Pten^WT^*, *Six3^Cre^;Pten^cHet^* and *Six3^Cre^;Pten^cKO^* retina cross-sections labeled by ChAT immunostaining show abnormal SAC somal lamination and disorganized dendrites in *Six3^Cre^; Pten^cKO^* retinas. **N.** Quantification of SAC dendrite stratification using IPLaminator shows aberrant dendrite stratification in *Six3^Cre^;Pten^cKO^* compared to controls (p = 0.0004). **O-Q.** P28 *ChAT^Cre^;Pten^WT^;Ai9*, *ChAT^Cre^;Pten^cHet^;Ai9* and *ChAT^Cre^;Pten^cKO^;Ai9* retina cross-sections with SAC somas and dendrites labeled via tdTomato. SAC somal lamination and dendrite organization are grossly normal, with two distinct bands in S2 and S4 in the IPL. **R.** Quantification of SAC dendrite stratification shows no significant changes in *ChAT^Cre^;Pten^cKO^;Ai9* SACs relative to controls (p = 0.4638). Data reported as mean ± SEM. Scalebars = 25 μm.

To examine how deletion of *Pten* affects SAC dendrite stratification in the IPL, we stained retinal cross-sections at P28. Similar to previous studies that examined pan-retinal deletion of *Pten*, we observed a gross disruption of SAC lamination, with highly disorganized S2 and S4 bands which appeared to bleed into S1 and S5 in *Six3^Cre^;Pten^cKO^* retinas (Figure 2K-N)[34, 35]. In contrast, we observed two well-defined tdTomato^+^ bands corresponding to S2 and S4 in *ChAT^Cre^;Pten^cKO^;Ai9* retinas (Figure 2O-R). While these bands appeared slightly less compact in *ChAT^Cre^; Pten^cKO^;Ai9* retinas compared to littermate controls, there was no statistical difference between genotypes when quantified. Therefore, PTEN is not required for SAC dendrite stratification in the IPL.

### Loss of *Pten* causes increased dendritic branching in SACs

SACs have a high degree of dendritic overlap with their neighbors, preventing analysis of individual cell dendrites at a population level. To perform comprehensive morphometric analysis of individual SACs, we induced sparse labeling with a *Cre*-dependent AAV (*AAV8-FLEx-tdTomato-CAAX*) injected into the vitreous of the eye at P1-P2. We analyzed *ChAT^Cre^;Pten^WT^*, *ChAT^Cre^;Pten^cHet^,* and *ChAT^Cre^;Pten^cKO^* SACs from both the GCL and INL at P21 when dendrite morphology is largely mature (Figure 3A-C”) [23]. We quantified total dendritic length, number of branch points, dendritic field area, and dendritic self-crossings (Figure 3D-G). While we did not detect any differences between *ChAT^Cre^;Pten^WT^* and *ChAT^Cre^;Pten^cHet^* SACs, there were significant changes in *ChAT^Cre^;Pten^cKO^* SACs. There was a small increase in total dendritic length in GCL SACs but not INL SACs in *ChAT^Cre^;Pten^cKO^* retinas (Figure 3D). The total number of branch points was significantly increased in both INL and GCL SACs in *ChAT^Cre^;Pten^cKO^* retinas, nearly doubling in number (Figure 3E). Despite the increase in dendrite branching, *ChAT^Cre^;Pten^cKO^* SACs show no changes in their dendritic field size (Figure 3F). We also found that *ChAT^Cre^;Pten^cKO^* SACs have a significant increase in dendritic self-crossings compared to control SACs (Figure 3G). Using a Sholl analysis to measure local changes in dendritic density we found that *ChAT^Cre^;Pten^cKO^* GCL SACs showed relatively localized increases in dendritic density in the distal 50% of their dendritic arbor (Figure 3H), whereas *ChAT^Cre^;Pten^cKO^* INL SACs showed a generalized increase in dendritic density across their entire arbor (Figure 3I). These results show that while PTEN is not required for establishing arbor size in SACs, it regulates proper dendrite branching. These local changes in density are significant as SACs are purely dendritic neurons with spatially segregated synaptic inputs and outputs [41, 42]. Therefore, local increases in dendritic density could lead to a biased recruitment of specific pre- and post-synaptic partners.

**Figure 3.**
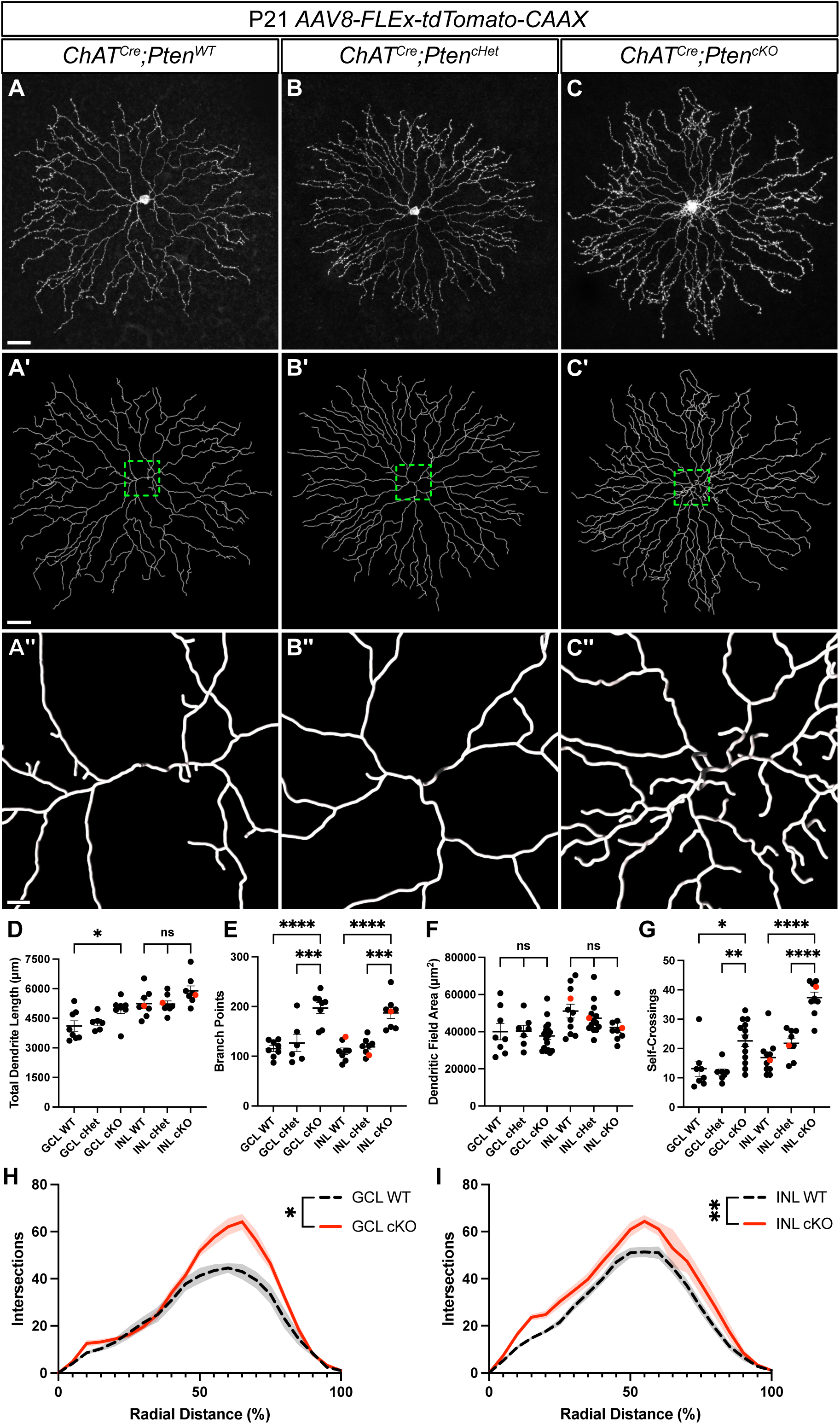
*Pten*-deficient SACs have abnormal dendritic branching patterns. **A-C**. SACs from P21 *ChAT^Cre^;Pten^WT^*, *ChAT^Cre^;Pten^cHet^*, and *ChAT^Cre^;Pten^cKO^* retina flat mounts sparsely labeled with *AAV8-FLEx-tdTomato-CAAX*. Images show single SACs located in the INL. **A’-C’.** Imaris reconstructions of SACs in A-C. **A’’-C’’.** Zoomed-in view of the dendritic arbor reconstruction near the soma. **D-G.** Quantification of total dendrite length (GCL p = 0.033; INL p = 0.076), number of branch points (GCL and INL p < 0.0001), dendritic field area (GCL p = 0.756; INL p = 0.152), and dendrite branch self-crossings (GCL p = 0.0015; INL p < 0.0001). A significant increase in the number of branch points and self-crossings is present in both GCL and INL SACs in *ChAT^Cre^;Pten^cKO^* retinas. Red dots indicate data from representative images. **H, I.** Sholl analysis reveals differences in local density that differ between GCL and INL SACs in *ChAT^Cre^;Pten^cKO^* retinas (GCL p = 0.0312; INL p = 0.0011). GCL cKO SACs show increased density near their terminal arbors, while INL cKO SACs show increased density throughout their arbor. Data reported as mean ± SEM and contain cells from at least 3 animals. Scalebars in A-C and A’-C’ = 25 μm. Scalebar in A"-C" = 3 μm.

### Alterations in dendrite branching arise late in the development of *ChAT^Cre^;Pten^cKO^* SACs and persist into adulthood

SACs undergo extensive dendritic arborization during the first two postnatal weeks, increasing their arbor territory and number of terminal branches [43]. To address when dendritic branching alterations arise in *ChAT^Cre^;Pten^cKO^* SACs, we selected two developmental time points, one prior to eye opening (P7) and one after (P14), to assess SAC morphology. To label P7 SACs, we used a genetic approach, crossing the *ChAT^Cre^;Pten* line with a *TIGRE-MORF* (*Ai166*) reporter line, which stochastically expresses EGFP in 1-5% of *Cre* positive cells [44]. At P7, SAC dendrites are in a highly dynamic state, constantly extending and retracting branches, which is critical for establishing their stereotyped radially symmetric morphology [43]. Since there were no differences between *ChAT^Cre^;Pten^WT^* and *ChAT^Cre^;Pten^cHet^* SACs at P21, we opted to include both genotypes as controls (*ChAT^Cre^;Pten^Ctrl^*). We focused our analysis on GCL SACs, as the imaging resolution of individual cells was better than INL SACs. Reconstruction and quantification of SACs identified no significant changes in total dendrite length, number of branch points, dendritic field area, or soma size between *ChAT^Cre^;Pten^Ctrl^* and *ChAT^Cre^;Pten^cKO^* SACs at P7 (Figure S1A-F). By P14, SACs have a much sparser dendritic arbor, with a morphology that nearly recapitulates their mature morphology. Using our sparse viral labeling approach to quantify morphology at P14, we found that total dendritic length, branch number, and dendritic field area remain unchanged in *ChAT^Cre^;Pten^cKO^* SACs compared to littermate controls (Figure S1G-K). However, we did detect a significant increase in soma size in *ChAT^Cre^;Pten^cKO^* SACs, suggesting that somal hypertrophy precedes changes in the dendritic arbor (Figure S1L). Taken together, our results indicate that loss of PTEN from SACs drives excess dendritic branching between P14 and P21.

To assess whether the increased dendritic branching seen at P21 would resolve, persist, or worsen in adulthood we injected *AAV8-FLEx-tdTomato-CAAX* at P28 and examined sparsely labeled *ChAT^Cre^;Pten^Ctrl^* and *ChAT^Cre^;Pten^cKO^* SACs at P60 (Figure 4A-B). Similar to SACs at P21, adult *ChAT^Cre^;Pten^cKO^* SACs showed a near doubling of dendrite branching across their arbor despite no change in total dendrite length (Figure 4D, E). Sholl analysis at P60 largely recapitulated the phenotypes at P21 as well, showing increased branch density in the outer 50% of the dendritic arbor (Figure 4G). However, we did identify distinctions between P21 and P60; notably, *ChAT^Cre^;Pten^Ctrl^* SACs had 3-5 proximal dendrites of similar sizes, whereas *ChAT^Cre^;Pten^cKO^* SACs frequently had a prominent single hypertrophic dendrite (Figure 4A’-B’). We defined any dendrite >1μm in caliber as a "hypertrophic dendrite" and found that these were present in 13/16 of *ChAT^Cre^; Pten^cKO^* SACs, compared with 1/12 in controls (Figure 4C). *ChAT^Cre^;Pten^cKO^* SACs also had slightly smaller dendritic field areas compared to *ChAT^Cre^;Pten^Ctrl^* SACs (Figure 4F). Despite these changes, SACs at P60 showed no changes in cell density, indicating that cell death was not occurring (Figure S2A-C). These results show that the long-term loss of *Pten* in SACs results in a persistent alteration in their dendritic arbor morphology.

**Figure 4.**
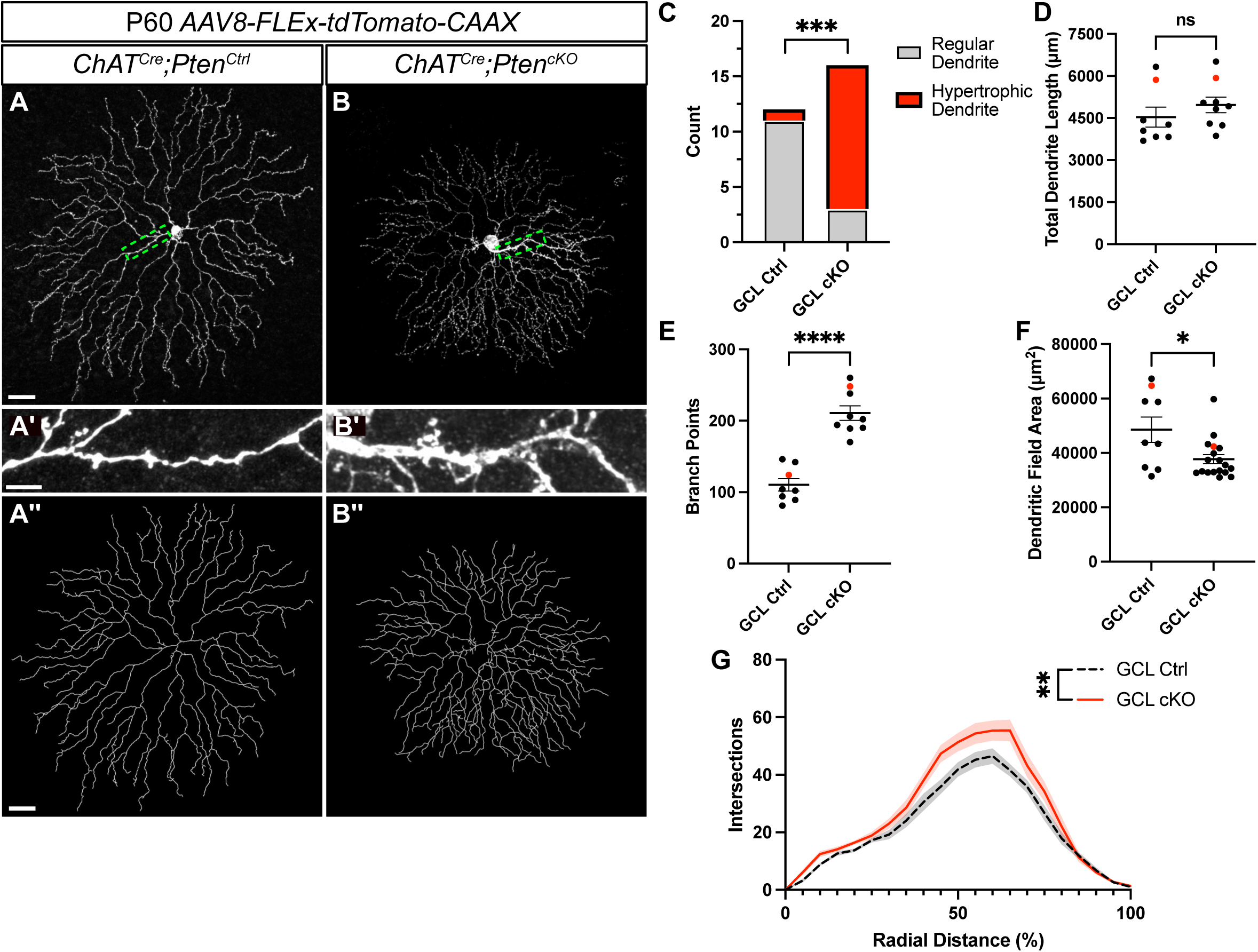
*Pten*-deficient SACs continue to show dendritic abnormalities at P60. **A-B**. P60 *ChAT^Cre^;Pten^Ctrl^* and *ChAT^Cre^;Pten^cKO^* SACs sparsely labeled by injection of *AAV8-FLEx-tdTomato-CAAX*. Images show single SACs located in the GCL. **A’-B’.** Enlargement from A-B highlighting that in *ChAT^Cre^;Pten^cKO^* SACs one of the dendrites frequently becomes hypertrophic. **A’’-B’’.** Imaris reconstructions of SACs in A-B. **C.** Quantification of the number of SACs containing a hypertrophic dendrite greater than 1μm in caliber (8.33% in *Pten^Ctrl^* and 81.25% in *Pten^cKO^* SACs; p = 0.0003 by Fisher’s exact test). **D-F.** Quantification shows that P60 cKO SACs have normal total dendritic length (p = 0.345), an increased number of branch points (p < 0.0001), and reduced dendritic field area (p = 0.013). Red dots indicate data from representative images. **G.** Sholl analysis reveals increases in local dendrite branch density similar to the phenotype observed at P21. (p = 0.0043). Data reported as mean ± SEM and contain cells from at least 3 animals. Scalebars for A and A” = 25 μm. Scalebars for A’ = 2 μm.

### Loss of *Pten* in SACs results in increased mTOR signaling over the course of development

PTEN serves as the primary negative regulator of the PI3K-AKT signaling pathway, which in turn activates mTOR signaling and inhibits GSK3β signaling (Figure 5K) [26]. Both mTOR and GSK3β alter the growth capacities of neurons and are likely candidates to regulate SAC branching [45]. We therefore examined how the loss of *Pten* from SACs affects these pathways using an antibody to pS6 as a readout of mTOR activity [35] and a genetically-encoded β-catenin:GFP reporter (*TCF/Lef:H2B-GFP*) as a proxy for GSK3β signaling (Figure 5A-H) [46]. In the GCL of P28 *ChAT^Cre^;Pten^cHet^* retinal flat mounts, pS6 was undetectable in SACs, while it was present in a subset of RGCs. In contrast, all GCL SACs in P28 *ChAT^Cre^;Pten^cKO^* retinas showed elevated pS6 levels, indicating activation of mTOR signaling (Figure 5I). Quantification of GFP signal in *ChAT^Cre^;Pten^cHet^*;*TCF/Lef:H2B-GFP* and *ChAT^Cre^; Pten^cKO^*;*TCF/Lef:H2B-GFP* retinas showed minimal fluorescence in SACs in both genotypes, suggesting that GSK3β signaling is unaffected by the absence of *Pten* (Figure 5J). Together, these results suggest that the morphological changes in *ChAT^Cre^;Pten^cKO^* SACs arise at least in part due to increased mTOR activity (Figure 5K-L).

**Figure 5.**
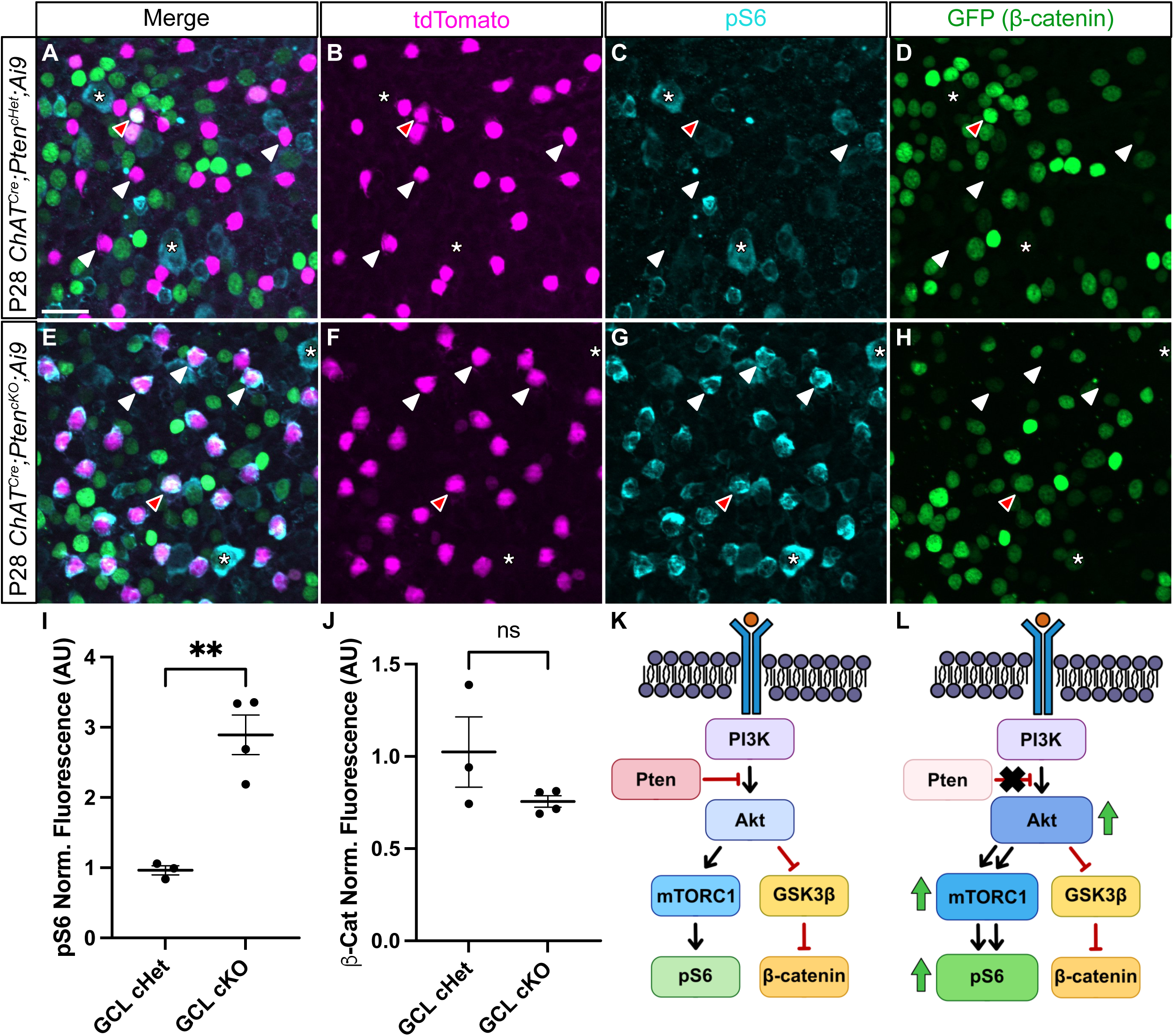
Deletion of *Pten* from SACs upregulates mTOR but not GSK3β signaling. **A-H**. Flat mount preparations of P28 *ChAT^Cre^;Pten^cHet^*;*Ai9;Tcf/Lef:H2B-GFP* and *ChAT^Cre^;Pten^cKO^*;*Ai9;Tcf/Lef:H2B-GFP* retinas immunostained for tdTomato (magenta, B, F), pS6 (teal, C-G), and β-catenin reporter *Tcf/Lef:H2B-GFP* (green, D-H). White arrowheads highlight SAC cell bodies in the GCL in both *Pten* controls (A-D) and cKOs (E-H). Red arrowhead indicates a SAC with elevated levels of β-catenin reporter signal. Asterisks indicate retinal ganglion cells in both the control and cKO retinas that show elevated levels of pS6. **I, J.** Quantification of pS6 and β-catenin fluorescence intensity show a significant increase in pS6 levels (p = 0.009) but no change in β-catenin levels (p = 0.312) in cKO SACs. **K, L.** Schematic showing a simplified view of the PI3K-AKT pathway. In the absence of *Pten* in SACs, AKT appears to increase mTOR activity as measured by pS6 levels, while GSK3β signaling as measured by β-catenin activity remains unchanged. Data reported as mean ± SEM. Scalebars = 25 μm.

Since SACs did not show any changes in dendritic branching during the most dynamic time of dendritic growth (P7-P14), we assessed pS6 at these ages in *ChAT^Cre^;Pten^cKO^* SACs. At P7, *ChAT^Cre^;Pten^cHet^* SACs showed high levels of pS6 immunoreactivity, which was not further elevated in *ChAT^Cre^;Pten^cKO^* SACs (Figure 6A-B, G, H). By P14 most SACs in *ChAT^Cre^;Pten^cHet^* mice had pS6 levels that were barely above background (Figure 6C-C’, G, H), whereas pS6 was significantly increased in *ChAT^Cre^;Pten^cKO^* SACs (Figure 6D-D’, G, H). Elevated pS6 levels were maintained in *ChAT^Cre^;Pten^cKO^* SACs at P60 (Figure 6E-F’, G, H). These results suggest that the lack of a dendritic branching phenotype in *ChAT^Cre^;Pten^cKO^* SACs at P7 may be because mTOR is already elevated at this age and loss of PTEN cannot drive further mTOR activity. In contrast, from P14 onwards mTOR activity has decreased in *ChAT^Cre^;Pten^cHet^* SACs, and deletion of *Pten* results in persistently elevated mTOR activity which maintains the branching and arborization process, leading to an increase in branch number by P21 and dendrite caliber by P60 (Figure 6I).

**Figure 6.**
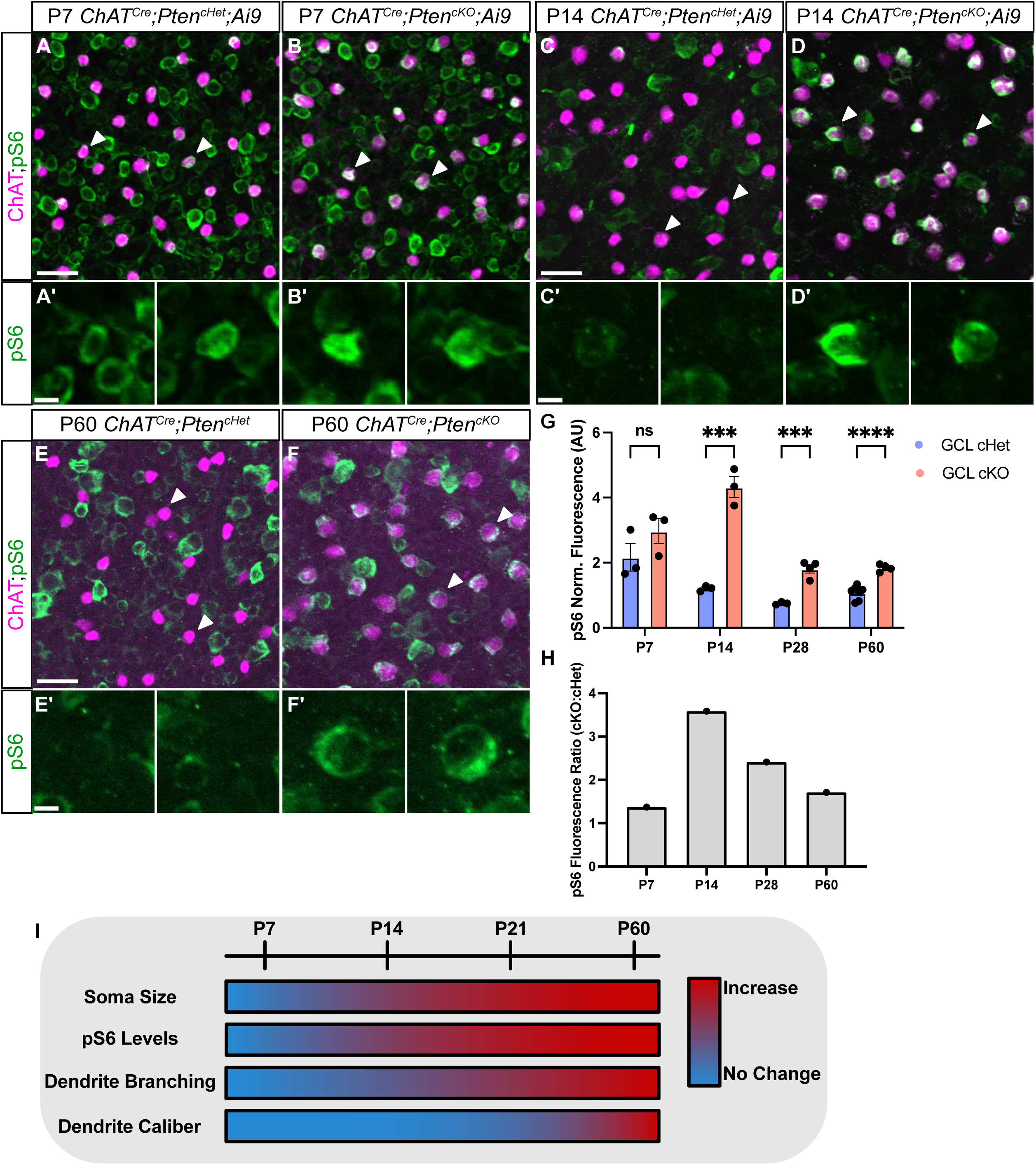
Increased pS6 precedes dendrite branching phenotypes in developing SACs. **A-F**. Retinal flat mounts of *ChAT^Cre^;Pten^cHet^* and *ChAT^Cre^;Pten^cKO^* SACs immunostained for ChAT (magenta) and pS6 (green) at P7 (**A-B**), P14 (C-D), and P60 (**E-F**). White arrowheads indicate SAC somas. **A’-B’.** Closeups of indicated P7 SAC somas show high levels of pS6 in both *ChAT^Cre^;Pten^cHet^* and *ChAT^Cre^;Pten^cKO^* retinas. **C’-D’.** At P14, pS6 levels are diminished in *ChAT^Cre^;Pten^cHet^* SACs but elevated in *ChAT^Cre^;Pten^cKO^* SACs. **E’-F’.** Closeups of SAC somas show that pS6 remains elevated in *ChAT^Cre^;Pten^cKO^* SACs relative to controls at P60. **G-H.** Quantification at P7, P14, P28, and P60 shows that pS6 levels are initially high in SACs at P7 (p = 0.2346) and decrease at later time points in control SACs, while pS6 levels remain significantly elevated in *ChAT^Cre^;Pten^cKO^* SACs relative to controls at P14 (p = 0.0007), P28 (p = 0.0006), and P60 (p < 0.0001). **I.** Summary of cellular phenotypes in *ChAT^Cre^;Pten^cKO^* SACs over the course of development and maturation. Increases in soma size and pS6 levels become apparent by P14, increased dendritic branching by P21, and localized increases in dendrite caliber are seen at P60. Scalebars = 25 μm in A, C, and E. Scalebars = 5 μm in A’, C’, and E’.

### SAC synaptic outputs and direction-selective circuit function are unaffected by loss of *Pten*

SACs have a highly compartmentalized synaptic organization, with presynaptic inputs from bipolar cells localized to the inner two thirds of their dendritic arbor, and their synaptic outputs localized to the outer third [47]. To examine whether the loss of *Pten* affected the number or compartmentalization of SAC synapses we intravitreally injected *AAV1-FLEx-mGFP-2A-Synaptophysin-mRuby* at P2 [48]. SACs transduced with this construct have a membrane-bound GFP that labels their dendritic arbor and synaptophysin (Syp) fused to mRuby to label synaptic outputs [49] (Figure 7A-B). Both *ChAT^Cre^;Pten^cHet^* and *ChAT^Cre^;Pten^cKO^* SACs showed robust localization of Syp:mRuby to the outer third of their dendritic arbor at P28 (Figure 7A’-B’). We quantified the number, volume, and spatial distribution of Syp:mRuby puncta and saw no differences between *ChAT^Cre^;Pten^cKO^* SACs and controls (Figure 7C-F). Therefore, even though SAC dendrite branching is dysregulated by P28 in *ChAT^Cre^;Pten^cKO^* SACs, synaptic outputs appear largely intact.

**Figure 7:**
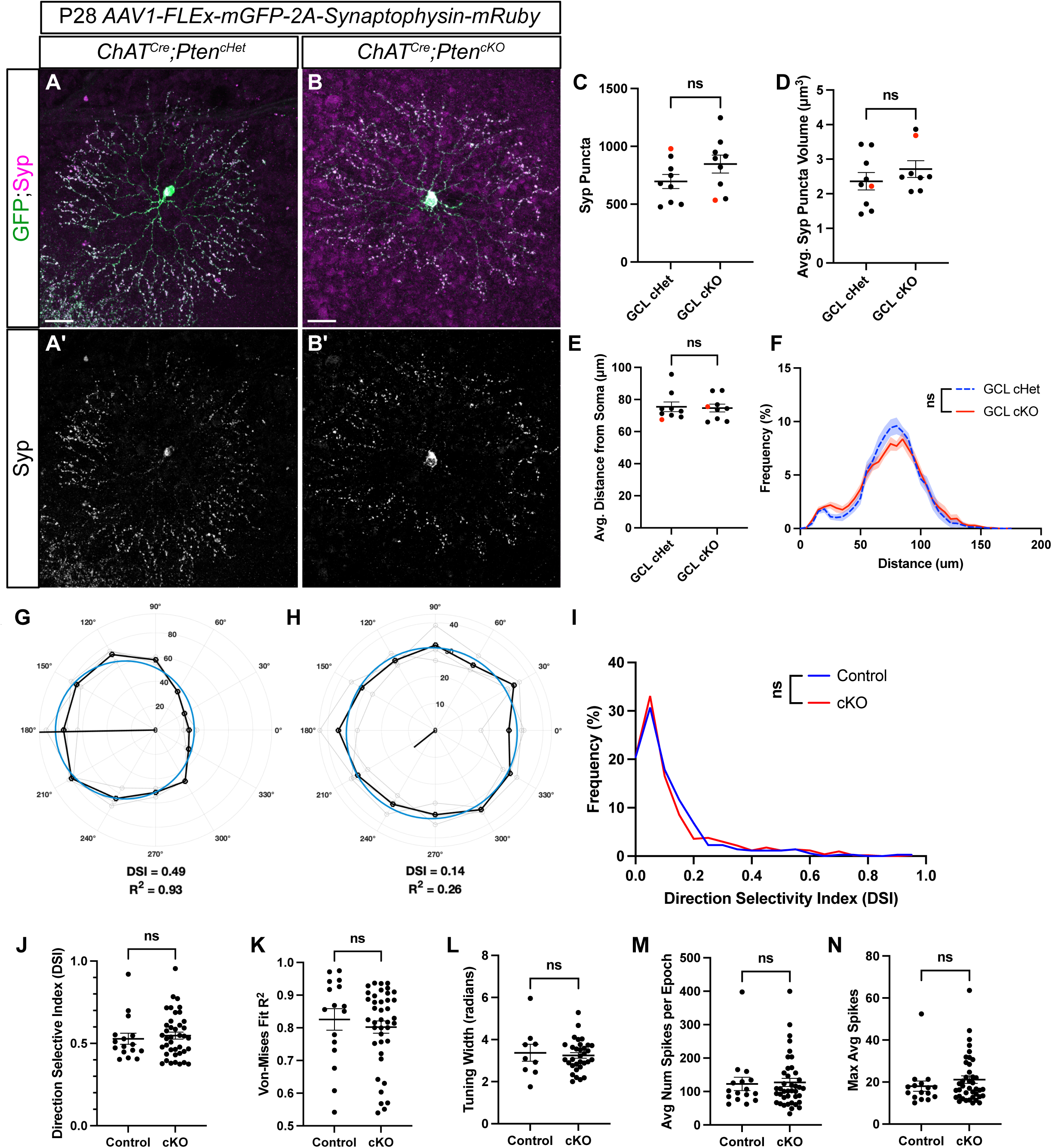
Loss of *Pten* does not affect SAC synaptic outputs nor alter retinal responses to directional stimuli. **A-B**. P28 *ChAT^Cre^;Pten^Ctrl^* and *ChAT^Cre^;Pten^cKO^* retinas injected with *AAV1-FLEx-mGFP-2A-Synaptophysin-mRuby* to label SAC dendrites with membrane-bound GFP and synaptic release sites with Synaptophysin (Syp) fused to mRuby. **A’-B’.** Syp:mRuby shows highly compartmentalized localization to the outer third of SAC dendritic arbors in both *ChAT^Cre^;Pten^cHet^* and *ChAT^Cre^ Pten^cKO^* SACs. **C-D.** Quantification of the number and volume of Syp:mRuby puncta show no significant differences between *ChAT^Cre^;Pten^cHet^* and *ChAT^Cre^ Pten^cKO^* SACs (p = 0.146 and p = 0.332). **E-F.** Quantification of the distribution of Syp:mRuby puncta reveals no significant changes in the average distance from the soma (p = 0.838) or the distribution along SAC dendrites (p > 0.999) in *ChAT^Cre^;Pten^cHet^* and *ChAT^Cre^ Pten^cKO^* SACs. Data reported as mean ± SEM and contain cells from at least 3 animals. Scalebars = 25 μm. **G, H.** Example polar plots of directional responses from individual RGCs following MEA recording of *ChAT^Cre^;Pten^cHet^* and *ChAT^Cre^ Pten^cKO^* retinas. Direction selective (DS) cells were identified based on their direction selective index (DSI) and their goodness of fit for the von Mises distribution. The black trace indicates the cells responses to light stimuli in different directions, while the blue circle represents the von Mises fit. **(G)** shows an example of a direction selective response, while **(H)** shows a direction insensitive response. **I.** Distribution of the DSI of all detected cells from MEA recordings shows that most cells fall below the DSI threshold for a DS cell (0.37), but a small population are direction selective (p = 0.9999). **J-N.** Quantification of DSI (p = 0.632), von Mises fit (p = 0.5179), tuning width (p = 0.6965), average number of spikes per epoch (p = 0.8411), and average spikes (p = 0.3397) in preferred direction from DS cells; no significant differences in the response properties of DS cells were detected between *ChAT^Cre^;Pten^cHet^* and *ChAT^Cre^ Pten^cKO^* retinas.

SACs have a conserved and well-characterized function of providing GABAergic inhibition and cholinergic excitation onto direction-selective ganglion cells (DSGCs) to tune direction selectivity [50, 51]. Genetic deletion of either *Sema6A* or *γ-Pcdhs* dramatically alters SAC morphology and degrades the direction selectivity of postsynaptic DSGCs [23, 52]. Therefore, we used multielectrode array (MEA) recordings to assess whether SAC-specific deletion of *Pten* influences downstream DSGCs. We isolated recordings from individual cells and computed their direction-selective index (DSI) in response to bars of light moving in 30-degree increments (Fig 7G-H). A von Mises goodness of fit test was performed to determine if a cell matched established DSGC response properties [53]. Cells with a DSI greater than 0.37 and a von Mises fit greater than 0.5 were classified as DSGCs. There was no significant difference in the distribution of DSIs of all RGCs between *ChAT^Cre^;Pten^cHet^* and *ChAT^Cre^;Pten^cKO^* retinas (Figure 7I). The average DSI, von-Mises fit, average number of spikes per epoch, average spikes in preferred direction, and tuning width in cells classified as DSGCs were unaffected in *ChAT^Cre^;Pten^cKO^* retinas compared to controls (Fig 7J-N). Taken together, these data show that while *Pten*-deficient SACs have significant morphological changes at P60, the function of the direction-selective circuit is unaffected.

## DISCUSSION

The process by which neurons develop stereotyped morphologies necessary to perform subtype-specific computations must be highly regulated to ensure consistency across each cell population. Here, we show that PTEN is required for SACs to adopt their precise branching patterns. By deleting *Pten* selectively from post-migratory SACs beginning at early postnatal ages, we were able to isolate PTEN’s cell-autonomous function and uncover its role in regulating dendrite branching at later stages of development. Mechanistically, this appears to result from elevated mTOR signaling, which normally decreases as SAC development progresses, but remains elevated in *Pten*-deficient SACs. Finally, we show despite altering the branching patterns of SAC dendrites, the loss of PTEN does not appear to disrupt the precise organization of synaptic outputs or the function of downstream retinal circuitry.

### Refining the cell-autonomous function of PTEN in SAC development

The environment in which cells develop plays a critical role in regulating their mature morphological features. Manipulating the pathways involved in regulating neuronal morphological development in intact preparations that maintain the extracellular environment can present challenges, as genetic deletions can lead to widespread anatomical changes that could affect neuronal morphology non-cell-autonomously. For example, pan-retinal *Pten* deletion broadly affects retinal progenitor proliferation, differentiation of multiple neuronal classes, somal lamination, and dendrite stratification of inner retinal neurons [34–36]. The SAC somal organization and dendrite stratification phenotypes in these *Pten*-deficient retinas resemble defects seen in mice lacking specific transmembrane adhesion proteins important for various aspects of SAC development, suggesting that PTEN could regulate the function of these proteins. Furthermore, deletion of *Pten* from retinal progenitors results in abnormal endocytic trafficking of cell surface proteins and signaling molecules important for SAC migration, mosaic spacing, and dendrite development [37]. However, it is equally possible that some of the SAC developmental phenotypes seen following pan-retinal *Pten* mutants are indirect and due to the profoundly disrupted extracellular environment surrounding them.

To address this issue, we compared the effects of pan-retinal (*Six3^Cre^;Pten^cKO^*) and SAC-specific (*ChAT^Cre^;Pten^cKO^*) deletion of *Pten* side-by-side to disentangle its non-cell and cell-autonomous roles in SAC development. Like prior studies, we observed profound disruptions in SAC somal lamination and dendrite stratification after pan-retinal deletion of *Pten*. In contrast, when this deletion was restricted to post-migratory SACs, we observed normal mosaic spacing and dendrite stratification in the IPL, suggesting PTEN is not required for these processes. This could be due to the timing of *Pten* deletion; pan-retinal deletion of *Pten* occurs in retinal progenitors between E8.5 and E9.5, whereas recombination in *ChAT^Cre^;Pten^cKO^* mice begins at P1 as these cells are tangentially migrating to space their somas and beginning to stratify their dendrites in the nascent IPL [20, 54–56]. However, mice in which *Megf10* or *PlexA2* are deleted using *ChAT^Cre^* still show defects in SAC mosaic arrangement and dendrite stratification, respectively [20, 23]. Therefore, the most likely explanation for our results is that PTEN is not essential for the function of the molecular pathways that regulate SAC somal positioning or dendrite stratification, while it is required for proper SAC dendrite branching. We also attempted to selectively delete *Pten* earlier from migrating SACs using *Megf10^Cre^* to examine SACs migration and mosaic spacing; however, these mice die immediately after birth, precluding these analyses.

### PTEN’s function in regulating neuronal morphology

PTEN has been highly studied in the nervous system due to its identification as an autism risk gene [57]. In cultured mammalian hippocampal neurons, knockdown of *Pten* increases dendrite branching through the PI3K-AKT-mTOR pathway [58]. *In vivo* deletion of *Pten* also causes generalized increases in dendritic growth and branching in hippocampal dentate granule cells, cortical neurons, and raphe serotonergic neurons [28, 29, 31, 33, 59, 60]. However, in *Drosophila* dorsocentral neurons, RNAi knockdown of *Pten* primarily caused localized branching as opposed to widespread neuronal hypertrophy, suggesting that PTEN’s role in regulating dendrite branching can differ depending on the neuronal subtype [61]. Similarly, we found while that *Pten*-deficient SACs displayed somal hypertrophy and nearly double the number of dendrite branches, they maintained their overall dendritic arbor size. It is unclear why deletion of *Pten* from SACs does not result in increased dendritic arbor size like it does in many other neuronal subtypes. Arbor size is tightly regulated in SACs, allowing them to create an even coverage factor of 30x across the retina. However, SACs do have the capacity to grow larger dendritic arbors, as deletion of the transmembrane protein *Amigo2* causes SAC dendrite length and overall dendritic arbor size to scale 50% larger, while branching is unaffected [25]. Based on the normal overall dendritic arbor size in *Pten*-deficient SACs, we can conclude that PTEN is not required for the regulation of dendritic length by AMIGO2. Instead, these findings reinforce that the molecular mechanisms that modulate SAC dendritic field size and the branching of their dendritic arbor are discrete processes. *Pten* deletion from SACs also did not significantly affect dendrite lamination in the IPL, suggesting it is not essential for the transmembrane proteins shown to regulate this process [22, 23].

Both γ-Pcdhs and Sema6A/PlexinA2 are required cell-autonomously within SACs to regulate dendrite self-avoidance [15, 23, 24, 43]. *γ-Pcdh* mutant SACs have a normal number of terminal branches and overall arbor size, while *Sema6A/PlexinA2* mutant SACs show reduced branching and arbor size. The dendrite morphology phenotypes that arise from mutations in these genes are much more severe that what we observed following deletion of *Pten*. Therefore, while it is possible that PTEN can function in the same molecular pathway as γ-Pcdhs and Sema6A/PlexinA2, there are clearly additional signaling pathways required. For PlexinA2, Rap1 GTPases are likely candidates, as SACs in mice with a point mutation that abolishes PlexinA2 RasGAP activity show significant dendritic self-avoidance phenotypes [24].

### Dysregulation of the PI3K-AKT-mTOR pathway following *Pten* deletion in SACs

Deletion of *Pten* from SACs is likely to have multiple effects, including dysregulation of the PI3K-AKT-mTOR pathway [26]. mTOR activation is a powerful enhancer of neuronal growth, and deletion of its upstream inhibitor *Tsc1* causes dendritic hypertrophy that largely recapitulates *Pten* deficiency in cortical and olfactory bulb neurons [45, 62]. Conversely, inhibition of mTOR via rapamycin is sufficient to rescue overgrowth phenotypes from *Pten* loss in dentate granule neurons [63]. Our results showing elevated pS6 levels following the deletion of *Pten* suggests that increased mTOR activity plays a major role in regulating dendrite branching in SACs. GSK3β, another effector downstream of the PI3-AKT pathway, can also regulate neurite outgrowth by modulating microtubule stabilization and polymerization [64, 65]. Our results using a genetic reporter of ý-catenin activity as readout of GSK3β function detected no differences between control and *Pten*-deficient SACs, suggesting that this pathway may not play a major role in regulating dendrite branching in SACs.

How does the loss of *Pten* in SACs result in increased dendritic branching? Elegant live imaging studies show that most of the SAC dendritic growth occurs between P4-P14 [43]. During the early portion of this phase, dendrites contain many exuberant self-contacting interstitial protrusions that are highly dynamic and largely prune away by P14. While we did not conduct live imaging in our studies, the timing of increased pS6 and increased branching provides some clues as to PTEN’s function. At P7 when SACs are undergoing extensive dendritic growth, high levels of pS6 were seen in control SACs. Deletion of *Pten* had no effect on pS6, somal hypertrophy, or dendrite branching at P7, suggesting that mTOR activity may already be near maximal levels at this age. By P14, we saw decreased levels of pS6 in control SACs, suggesting that mTOR activity normally declines coincident with when dendritic arbor growth and branching begins to slow. In contrast, *Pten*-deficient SACs show elevated pS6 and somal hypertrophy at this age, yet dendrite branching is unaffected. It is not until P21 and later that *Pten*-deficient SACs show increased dendrite branching. This suggests that loss of PTEN does not generally affect the major phase of developmental dendrite growth in SACs, but prolongs it beyond its normal plateau, resulting in excessive branching. The presence of excessive dendrite branches at P60 suggests that these branches are not unstable dynamic projections like those seen in developing SACs but rather are persistent branches. Whether these aberrant branches are functionally integrated into retinal circuitry is unclear.

At P60, SACs in *ChAT^Cre^;Pten^cKO^* retinas began to show a decrease in dendritic arbor size, suggesting that prolonged deletion of *Pten* could have adverse effects in neurons. In Purkinje neurons, loss of *Pten* results in the eventual apoptotic death of these neurons beginning around 6 months of age [30]. We did not observe any loss of SACs in *ChAT^Cre^;Pten^cKO^* retinas at P60, and did not examine later ages as these mice eventually develop facial tumors [66]. However, it is important to consider the long-term consequences of *Pten* deletion in neurons, as it is widely studied for its ability to facilitate axon regeneration after injury [67–70]. While these studies rarely report the effect of *Pten* deletion on the dendrites of these cells, a recent report shows that RGC dendrites rapidly retract in response to axonal injury, and co-deletion of *Pten* and *SOCS3* exacerbates this effect, although it is unclear whether these dendritic arbors eventually recover in size [71]. Our results suggest that the long-term deletion of *Pten* in neurons can cause deleterious changes in dendritic architecture.

### Functional Consequences of *Pten* Loss in SACs

At a circuit function level, SACs are crucial for the directional tuning of downstream DSGCs, as eliminating or pharmacologically silencing SACs results in a loss of direction selectivity [72–74]. SACs themselves display intrinsic direction selectivity, responding preferentially to centrifugal motion moving from the soma to the dendritic tips [75, 76]. Mutations that cause disruptions of the radial morphology of SAC dendritic arbors result in defective direction selectivity [23, 52, 77]. Loss of *Pten* in other neurons can dramatically affect their circuit function; hippocampal dentate granule neurons and serotonergic raphe neurons lacking *Pten* are hyperactive due to changes in their intrinsic excitability and have an increased number of excitatory inputs, whereas *Pten*-deficient Purkinje neurons show reduced excitability [28, 30, 32, 33, 78]. It is therefore somewhat surprising that despite the abnormal dendritic branching in SACs, direction selectively appeared intact in *ChAT^Cre^;Pten^cKO^* retinas. We did not examine the intrinsic physiological properties of *Pten*-deficient SACs directly, so it is possible that there are cell-autonomous functional differences that are not sufficient to affect the downstream DSGCs. We also observed no changes in the localization, density, or size of SAC synaptic outputs *ChAT^Cre^;Pten^cKO^* retinas. While the molecular mechanisms that underlie the spatial segregation of synaptic inputs and outputs in SACs remain unknown, *Pten* signaling is apparently not required.

Altogether, our study refines the role of PTEN in regulating the morphology of SACs, showing that it is critical for establishing the highly stereotyped dendritic branching pattern in these neurons. Further studies will be needed to identify additional intracellular signaling pathways that function downstream of cell surface receptors to regulate cytoskeletal dynamics in developing SACs.

## STAR★METHODS

### Resource Availability

#### Lead contact

Further information and requests for resources and reagents should be directed to and will be fulfilled by the Lead Contact, Dr. Kevin Wright

#### Materials Availability

Transgenic mouse lines used in this study are available upon request or in a central repository (Jackson Laboratory). All antibodies and reagents are commercially available.

#### Data availability

Data available upon request to lead contact.

### Experimental Models and Subject Details

#### Animals and Animal Procedures

All animal procedures were approved by the Oregon Health and Science Institutional Animal Care and Use Committee (IACUC). The following mouse lines were used: *ChAT^Cre^* [79], *Six3^Cre^* [80], *Pten^flox^* [81], *Ai9* [82], *TIGRE-MORF/Ai166* [44], *TCF/Lef:H2B/GFP* [46]. All lines were maintained on a *C57BL/6J* background. Mice of both sexes were used for experiments. Intravitreal injections were performed on mice at P2 or P28. P2 mice were anesthetized through indirect contact with ice and brought back to body temperature through contact with a warm glove. Mice were injected with a 30psi pulse for 30ms. P28 mice were anesthetized using isoflurane at a flow rate of 3%, and maintained at a flow rate of 1.5%. After deeply anesthetized, animals were placed on a Kopf stereotaxic injection rig (Model 1900 Stereotaxic Alignment system). 0.5% proparacaine hydrochloride ophthalmic solution was applied as a topical anesthetic to the eye, followed by 1% tropicamide ophthalmic solution for dilation. Gentle-eye lubricant was added to both eyes, and a micro vessel clamp was used to push the globe outwards. Needles were pulled on a Sutter Instrument micropipette puller (P-97 Flaming/Brown Micropipette Puller) and beveled with sandpaper. Animals were injected with *AAV8-FLEx-tdTomato-CAAX* at a titer of 2.6x10^11^ or *AAV1-FLEx-mGFP-2A-Synaptophysin-mRuby* at a titer of 1.8 x 10^12^ [48].

### Method Details

#### Tissue Processing and Immunohistochemistry

Mouse eyes were enucleated and drop fixed in 4% EM-grade PFA for 30 minutes. For *TIGRE-MORF* mice, retinas were instead fixed with a solution of 9% Glyoxal 8% Acetic Acid at pH = 4.0, as this improved resolution of the Tigre-GFP signal. After fixation eyes were washed in PBS. Eyes were then pierced with a 30G ½ inch needle and the cornea was cut away with microdissection scissors.

For retinal flat mounts, the retina was isolated and then transferred to an Eppendorf tube and blocked and permeabilized with a blocking solution (2% normal donkey serum, 0.2% Triton X-100, 0.002% Sodium Azide) for 1 hour. After blocking, retinas were stained with primary antibodies and left shaking at 4°C for 3-4 days. Retinas were then washed overnight in PBS. Retinas were then stained with secondary antibodies and left shaking at 4°C for 1 day. Four cuts were made into the retina to allow it to lay flat on glass microscope slides. Retinas were mounted with Fluoromount-G and sealed with nail polish. All antibody dilutions are listed on the Key Resource Table.

For retinal cross-sections, the retina and lens were left in the eye cup and cryoprotected in 15% sucrose overnight. The next day, the lens was removed, eyes were placed in Neg-50, and the eyes were frozen in the eye cup in 2-methylbutane. Eyes were cryosectioned on a Leica cryostat (CM3050 S) at 20μm sections. The edges of the slide were then coated with a hydrophobic barrier using an ImmEdge pen. The slide was washed with PBS to remove any remaining Neg-50 attached to the slide. Retinas were then stained with primary antibody at 4°C overnight. The next day, retinas were washed with PBS three times for 15 minutes. Secondary antibody was applied and the retinas were left at room temperature for 2 hours. Hoechst was then applied to the retinas for 10 minutes, followed by three 15-minute PBS washes. Retinas were mounted with Fluoromount-G and sealed with nail polish. Antibody dilutions were the same as in flat mounts.

#### Fluorescence Image Acquisition

Retinal sections were imaged on a Zeiss Axio Imager M2 upright microscope equipped with an ApoTome2 using a 20x objective. Retinal flat mounts imaged for cell density, somal quantification, mosaic analysis, pS6, and β-catenin quantification were also imaged with these settings. Retinal flat mounts imaged for single cell morphology and synaptophysin labeling were imaged on a Zeiss LSM 900 confocal microscope using a 40x water objective with NA = 1.2.

Images were acquired using the Zeiss Zen Imaging software for both microscopes.

#### Multi-Electrode Array Recordings

Tissue from mouse retinae were placed RGC side down on a 3Brain Accura HD-MEA connected to a BiocCAM DupleX recording system (3Brain AG, Wädenswil, Switzerland). The Accura HD-MEA contains 4096 electrodes in a 3.8 x 3.8 mm area, where each electrode is 21 µm x 21 µm spaced 60 µm apart. The internal diameter of the reservoir is 25mm, with a 7 mm height.

Retinae were dissected off the choroid and the vitreous was then carefully dissected from the tissue prior to mounting the tissue photoreceptor-side down on Millicell polytetrafluoroethylene membrane cell culture insert (Millipore Sigma; PICM0RG50). To adhere the retina to the membrane, the Ames medium was removed from the insert and gentle suction was applied using a gas stone connected to a vacuum chamber with filter paper between the stone and the membrane. The insert was then placed back into Ames’ medium and the membrane trimmed to outside the edges of the retinae. The retinae/insert was placed RGC-side down on the MEA surface in Ames’ medium and the Ames medium was removed from the reservoir to facilitate connectivity with the electrodes. A platinum harp was placed over the insert to hold the retinae in place, the reservoir was filled with Ames, the MEA was transferred to the BioCAM DupleX, and the reservoir was continuously superfused with Ames’ medium @ 32°C.

Visual stimuli were generated using custom software generated in Python (PyStim; https://github.com/SivyerLab/pystim) and presented on a LightCrafter 4500 projector (Texas Instruments, USA) modified by removing the focusing optics. Projector light was captured with a TV lens, passed through neutral density filters and an Olympus MVX10 fluorescence microscope system with a MVPLAPO 2XC (0.5 numerical aperture) objective. A 1.5 mm coverslip was mounted onto the reservoir to reduce diffraction caused by the air/Ames interface. Visual stimuli presented used either the green (LE CG Q9WP; 520 nm peak) or blue LEDs (LE B Q9WP; 455 nm peak) and were presented full field at a range of light intensities between 6.2e11 photons/cm2/s and 8.9e13 photons/cm2/s. Stimuli consisted of green full field chirp responses used to isolate rod and cone mediated inputs to RGCs, ON and OFF responses, and temporal and spatial frequency tuning properties. Moving stimuli consisted of gratings moving in 12 directions for 3 seconds each direction.

### Quantification and Statistical Analysis

#### SAC Population Analyses (Cell Counting, Soma Size, Mosaic Spacing, and pS6 and β-catenin)

Retina flat mounts were immunostained for either tdTomato or ChAT. Three regions were imaged per retina per animal. GCL and INL images were captured from the same regions of the retina, avoiding the center and far periphery. Every *Pten^cKO^* animal included had at least one *Pten^cHet^* littermate control. The tdTomato or ChAT channel was binarized in ImageJ using either the Otsu or Huang thresholding algorithm and ImageJ’s “Analyze Particles” function was used to perform cell counts. As part of this process, we generated individual regions of interest (ROIs) for every cell, an ROI consisting of every cell and an inverse ROI consisting of the background. As part of cell density quantification, the size of the particles and the xy coordinates of the center of SAC somas were collected for soma size and mosaic spacing analysis. These xy coordinates were then entered into WinDRP, a program designed to calculate the regularity of cells using a density recovery profile (Rodieck 1991). From this, a regularity index ratio was calculated, which defines how mosaically spaced a population of cells are compared a random distribution. The same image acquisition and ImageJ pipeline for cell counting was used to analyze pS6 and β-catenin fluorescence. ImageJ’s “Analyze Particles” function was used to obtain ROIs for the SACs within an image and the inverse ROI representing background signal. The pS6 and β-catenin fluorescent signals were then measured in both these ROIs. The SAC ROI was then normalized to the background ROI to obtain a normalized fluorescence intensity signal.

#### IPLaminator Analysis

Lamination was quantified in P28 retina cross-sections stained for tdTomato and DAPI using IPLaminator. IPLaminator is an ImageJ plugin designed to bin and quantify retinal lamination [83]. DAPI was used to define the area of the IPL, then tdTomato fluorescence was measured by IPLaminator. For every image, the IPL was divided into 20 equal sections and measured along the depth of the IPL to normalize any variance in IPL thickness.

#### Imaris Reconstructions and Morphometric Analysis

SACs were reconstructed manually using the Imaris Filaments module. Briefly, the soma was assigned as a dendrite beginning point and dendrites were traced from there using the AutoPath tool. From these reconstructions, data including total dendrite length, number of branch points, and Sholl intersections were collected. Sholl data was generated from Imaris at 1μm intervals. Sholl data was normalized to the radial distance of the cell by averaging the number of Sholl intersections along every 10% of the radial distance. Dendrite field area was measured in ImageJ by taking the convex hull of the fluorescent area covered by a SAC. The number of self-crossings at P21 was manually counted in ImageJ by counting dendrite branch intersections in single z-planes across a z-stack. Dendrite caliber was measure in ImageJ by drawing a line perpendicular to the dendrite and measuring its length. Any dendrite greater than 1μm in caliber was considered a hypertrophic dendrite.

#### Synaptophysin Analysis

Synaptophysin puncta were quantified in Imaris using the Surfaces module. Briefly, surfaces were generated for all putative synaptophysin puncta, which were then filtered based on fluorescence of the membrane bound GFP signal to exclude noise outside the SAC. Puncta smaller than 0.5μm were also excluded. The number of puncta, size of the puncta, and distance from the soma of every puncta was obtained through Imaris.

#### Spike Sorting

Herding Spikes 2 (HS2), via SpikeInterface^2^ Python framework, was used for spike detection and sorting. HS2 uses a mixed approach of spike spatial and prominent waveform features combined with a mean shift clustering algorithm to identify individual cells and their corresponding spikes on the array. The spike sorting scripts were executed on a high-performance computing cluster (exacloud).

The following HS2 parameters were used:

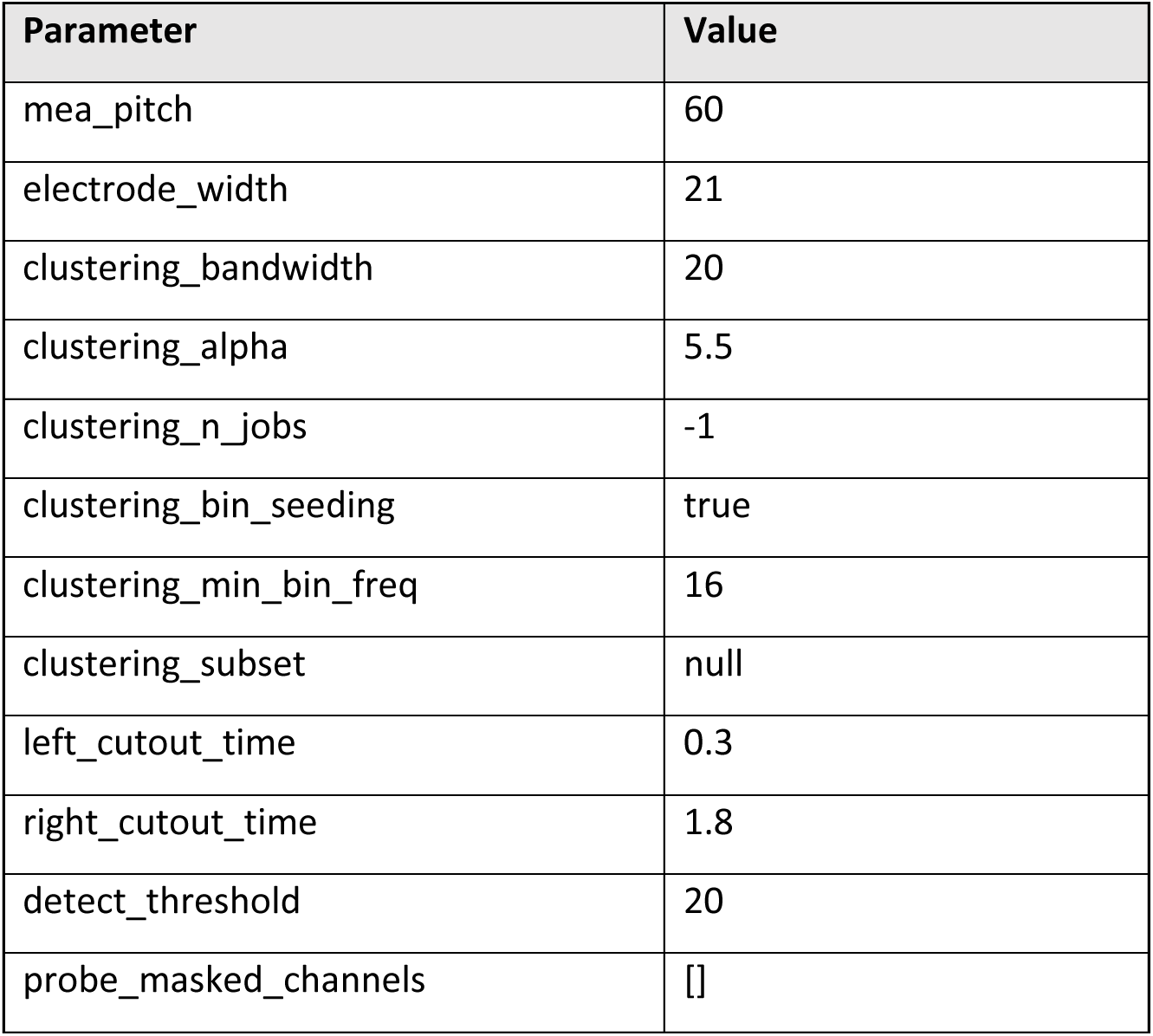
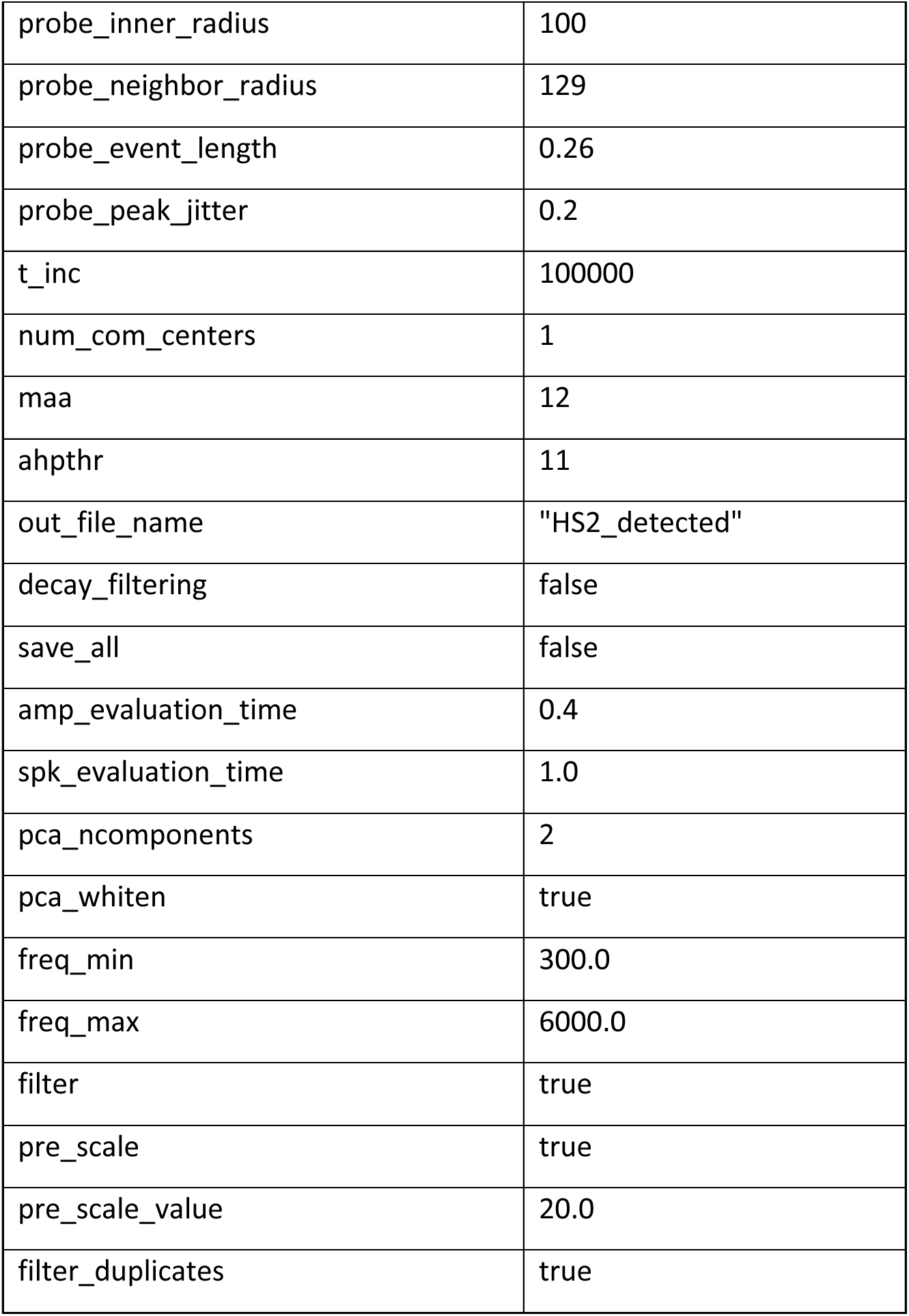

#### DSGC Classification and Statistical Analysis

Post spike sort, subsequent analysis in MATLAB R2023a (Mathworks) was performed for further cell classification based on light responses. Responses from individual units were assessed for each presented direction of light. Units without a minimum of 400 total spikes were filtered out. Remaining units had their direction selectivity 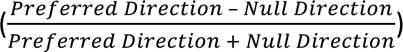 and von Mises fit calculated [53]. Units with a DSI greater than 0.37 and a von Mises fit greater than 0.5, were classified as putative DSGCs. Additional filtering was then performed to make sure the units had a minimum of 10 average spikes in their preferred direction across the epochs. Once DSGCs were determined, the average number of spikes per stimulus window (epoch) and the average number of spikes in their preferred direction were measured. Tuning width was calculated by using the full width at half maximum (FWHM) [53].

#### Statistics

For each experiment and time point, a minimum of three mice per condition were analyzed. For single cell morphological analysis, a minimum of 6 cells were analyzed, with most conditions having at least 8 cells. For all datasets, the variance was reported as mean ± SEM. For analysis between two groups, a Student’s t-test was performed. For analysis between three groups, an ANOVA with Tukey’s multiple comparison was performed. For comparisons of distributions, a Kruskal-Wallis test was performed. For Sholl Analysis, an area under the curve (AUC) analysis was performed, where AUC statistics (mean, SEM, n) were computed, then analyzed via a Student’s t-test. For comparison of the ratio of SACs with a hypertrophic dendrite, Fisher’s exact test was used. All statistical tests were performed in GraphPad Prism 9.

## Key Resources Table

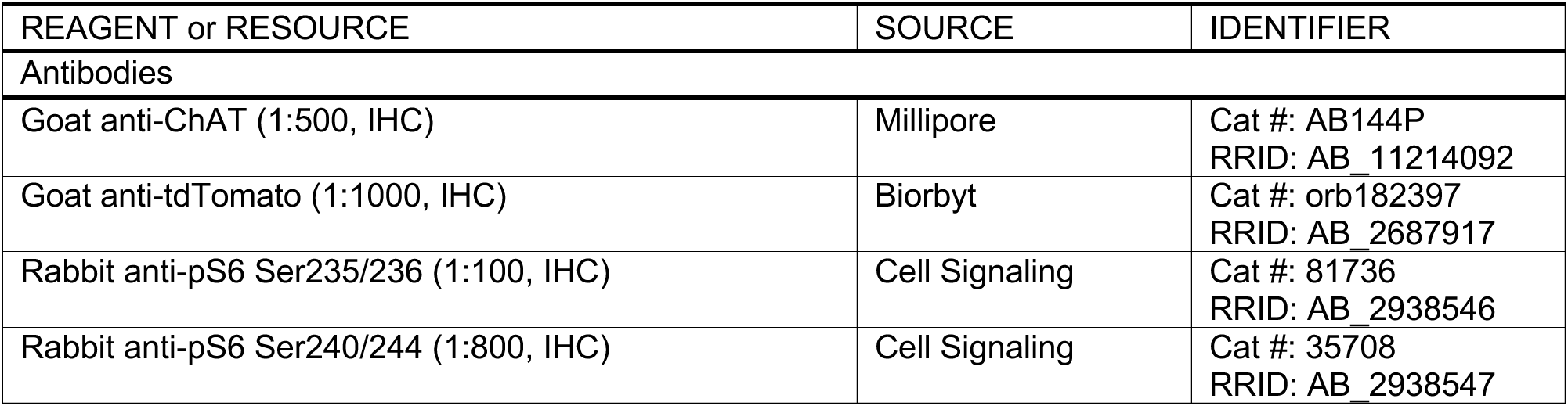
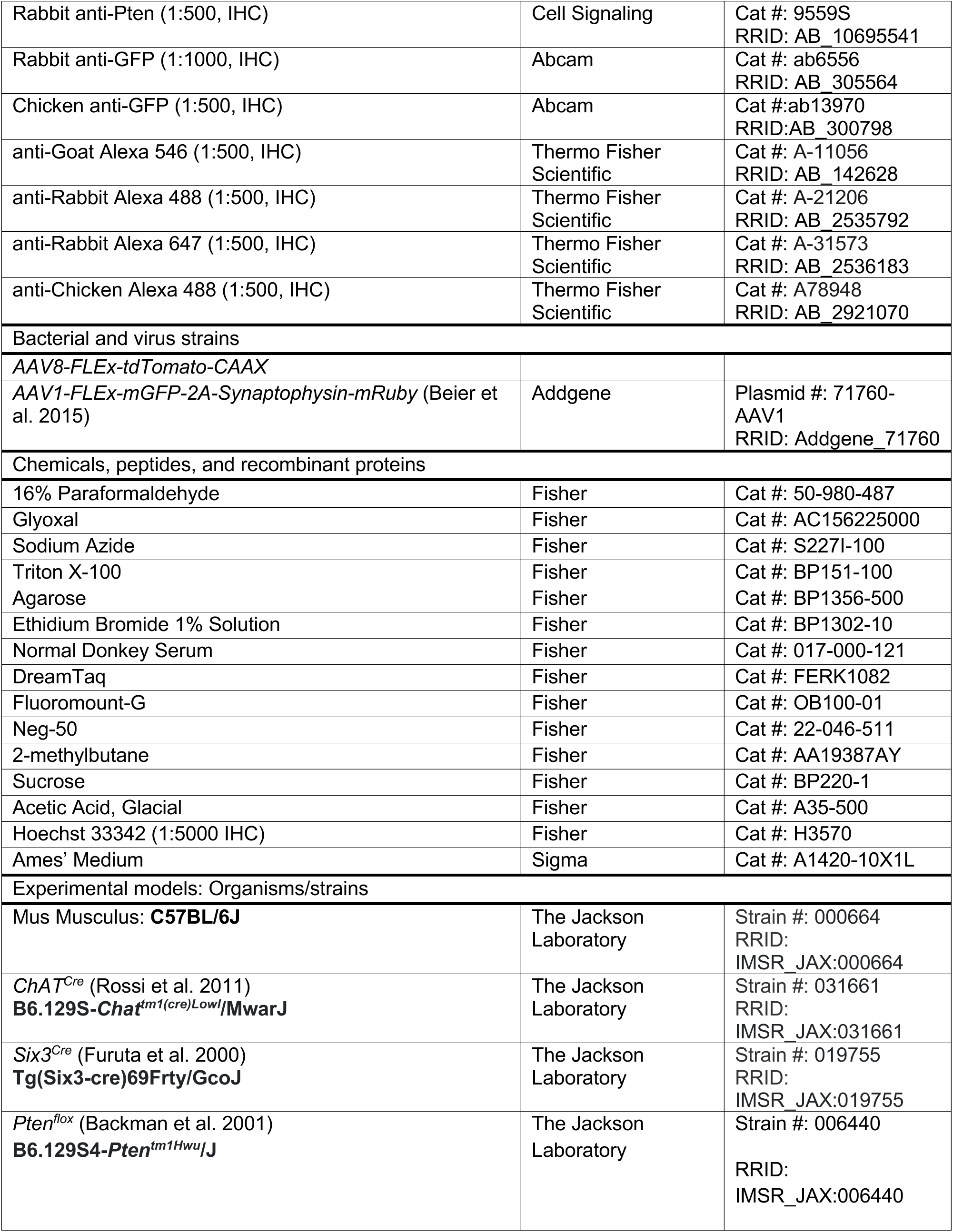
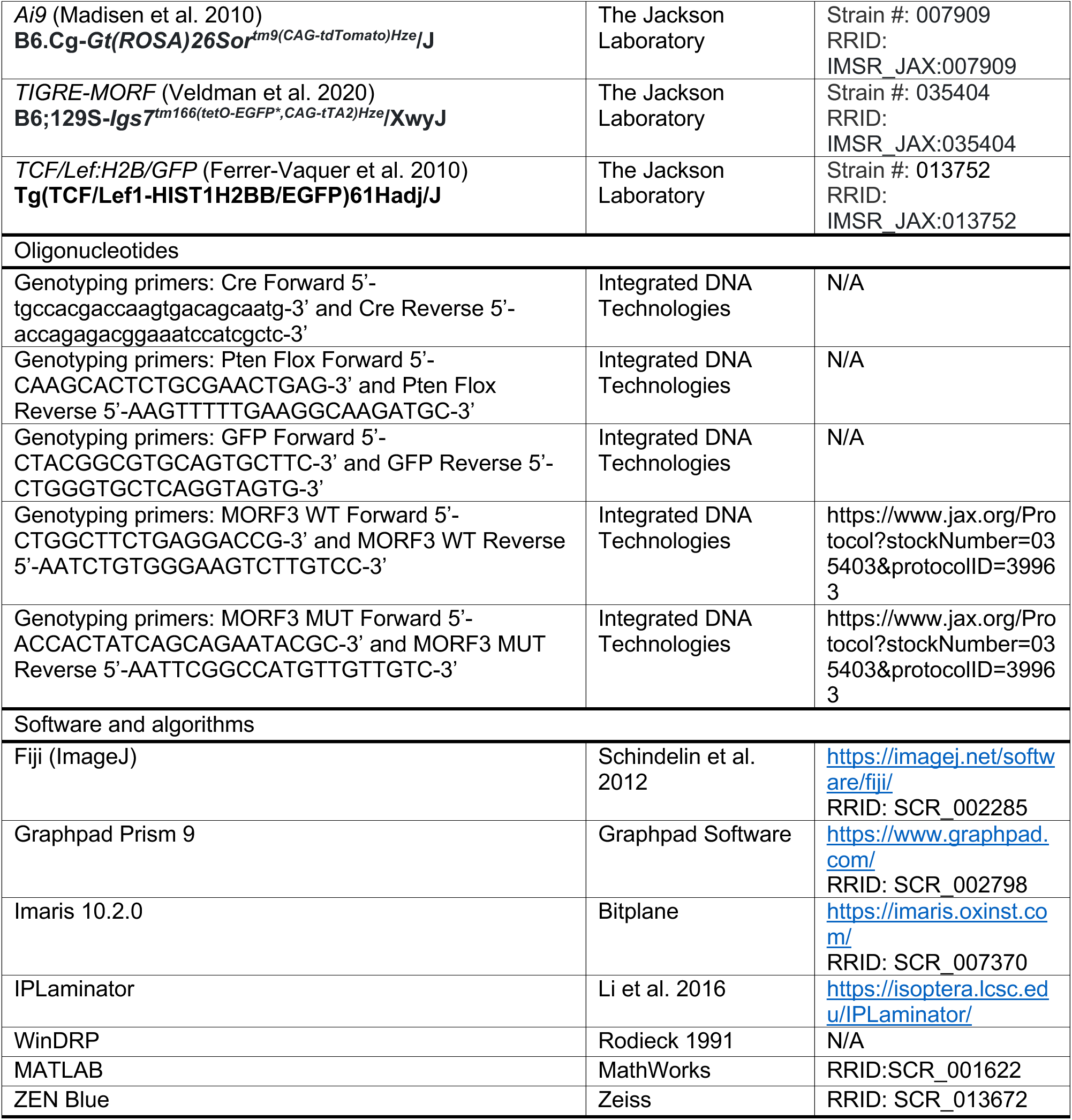

## Acknowledgments

We acknowledge expert technical assistance by staff in the OHSU Advanced Light Microscopy shared resource and OHSU Department of Comparative Medicine. We would like to thank members of the Wright laboratory for their comments on the manuscript. This work was funded by NIH Grants R01EY032057 (KMW), T32EY023211 (TB), P30NS061800 (OHSU ALM)

---

**Figure S1.**
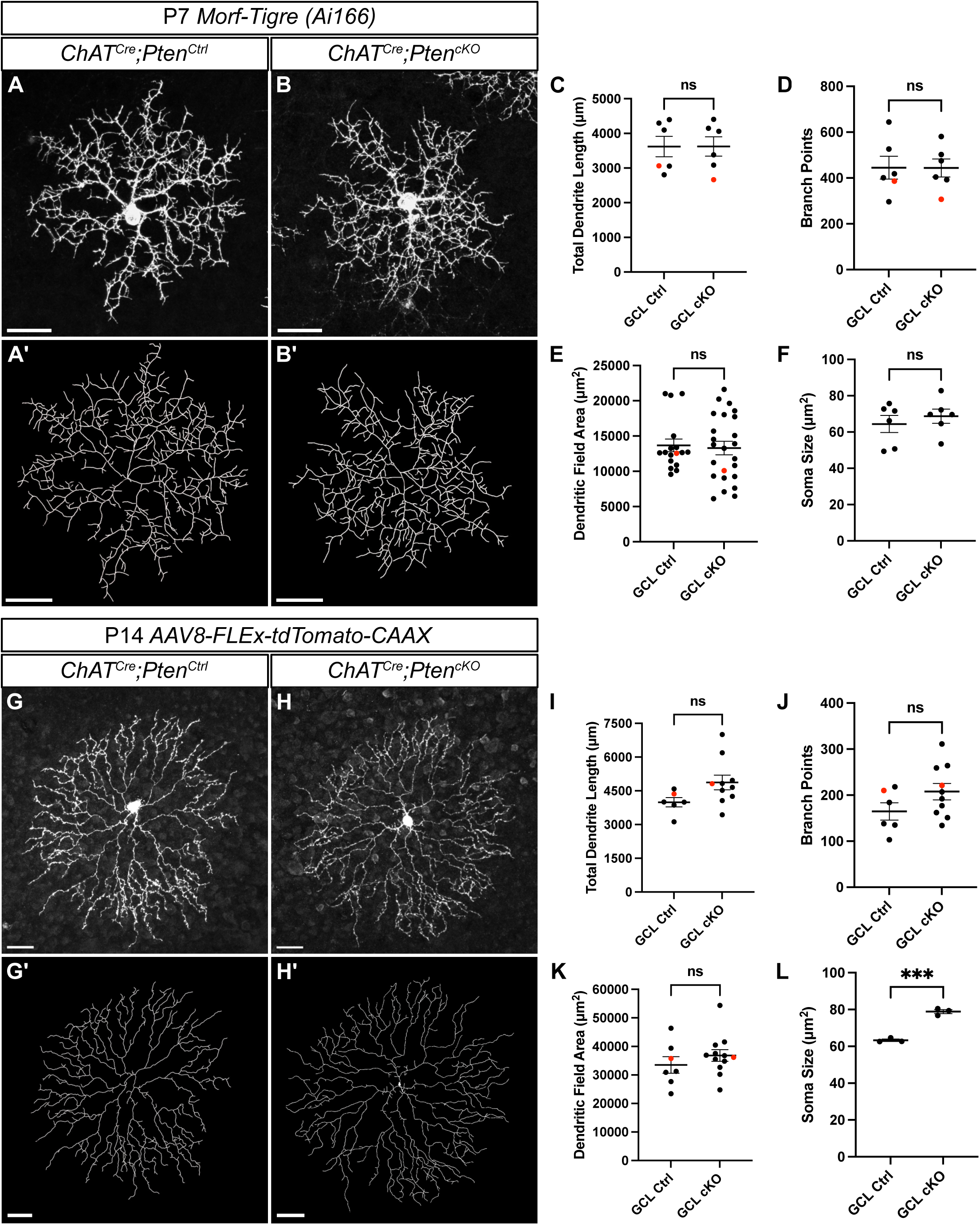
*Pten* deletion from SACs does not affect their morphology at early developmental timepoints. **A-B**. P7 SACs from *ChAT^Cre^;Pten^cHet^*;*Ai166* and *ChAT^Cre^;Pten^cKO^*;*Ai166* retinas were sparsely labeled using a genetically encoded *Morf-Tigre* reporter. Images show single SACs located in the GCL. **A’-B’.** Imaris reconstructions of P7 SACs from A-B. **C-E.** Quantification of total dendrite length (p = 0.778), number of branch points (p = 0.889), and dendritic field area (p = 0.784) from individual SACs showed no significant differences between control and cKO SACs at P7. **F.** Quantification of average soma size (p = 0.495) by animal showed no changes between control and cKO SACs. **G-H.** P14 *ChAT^Cre^;Pten^cHet^* and *ChAT^Cre^;Pten^cKO^* SACs were sparsely labeled by injection of *AAV8-FLEx-tdTomato-CAAX*. Images show single SACs located in the GCL. **G’-H’.** Imaris reconstructions of P14 SACs from G-H. **I-K.** Quantification of total dendritic length (p = 0.072), number of branch points (p = 0.137), dendritic field area (p = 0.361) from individual SACs showed no significant differences between control and cKO SACs. **L.** Quantification of average soma size revealed significant increases in cKO SACs at P14 (p = 0.0003). Red dots indicate data from representative images. Data reported as mean ± SEM and contain cells from at least 3 animals. Scalebars = 25 μm.

**Figure S2.**
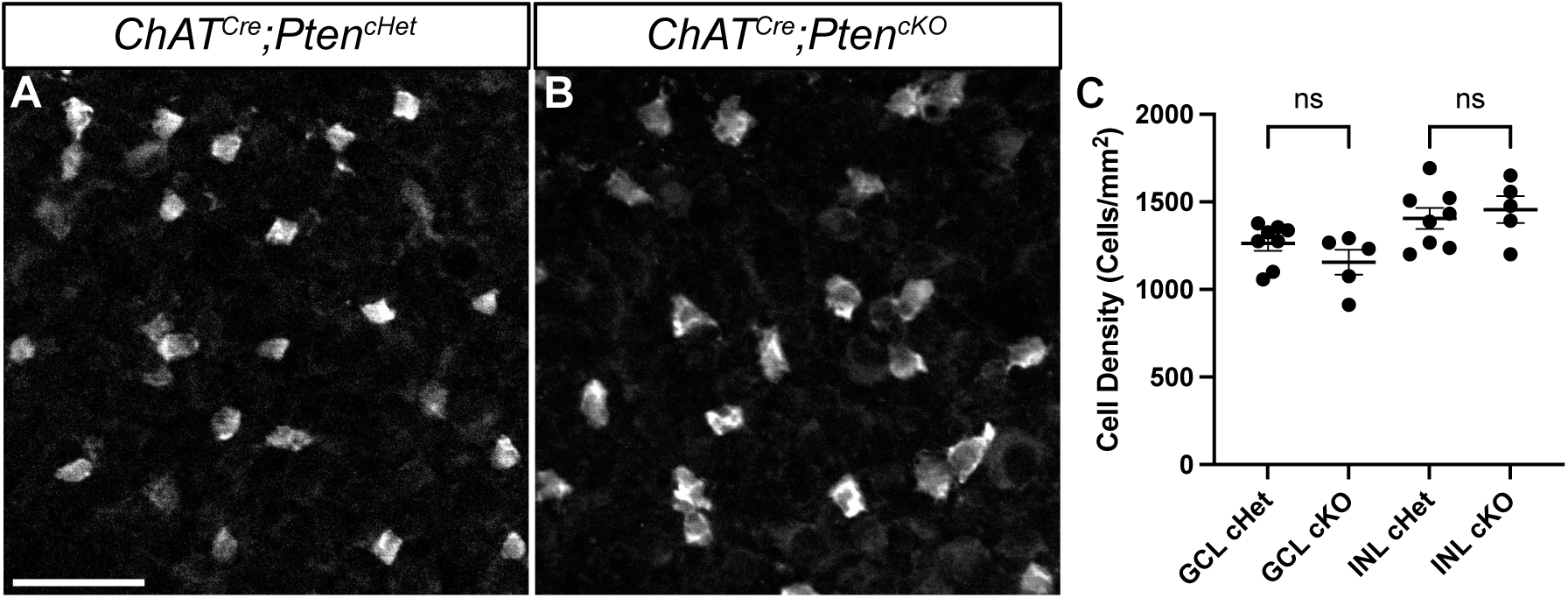
*ChAT^Cre^* mediated *Pten* deletion from SACs does not cause changes in cell density at P60. **A, B.** Representative images of P60 SACs from the GCL labeled with ChAT antibody. **C.** Quantification of cell density reveals no changes in cell density between *ChAT^Cre^;Pten^cHet^* and *ChAT^Cre^;Pten^cKO^* SACs at P60 in both the GCL (p = 0.1931) and INL (p = 0.6171). Data reported as mean ± SEM. Scalebars = 25 μm.

